# Long-distance dispersal drives global tropical distributions in a widespread moth lineage (Lepidoptera: Limacodidae)

**DOI:** 10.64898/2026.05.16.724310

**Authors:** Tabitha R. Taberer, Marianne Espeland, Sebastian Martin, Tim Coulson, Sonya M. Clegg

**Affiliations:** Department of Biology, University of Oxford, Life and Mind Building, Oxford, South Parks Road, OX1 3EL, UK; African Natural History Research Trust, Street Court, Kingsland, Leominster, HR6 9QA, UK; Leibniz Institute for the Analysis of Biodiversity Change, Museum Koenig, Adenauerallee 127, 53113 Bonn, Germany

**Keywords:** Global tropics, Limacodinae, Museum specimens, Pantropical, Whole genome sequencing

## Abstract

Understanding how global biodiversity patterns arise is a central theme of biogeography, with contemporary theory recognising the roles of both dispersal and vicariance. Genera that are broadly distributed can provide important systems for disentangling the relative influence of these processes across evolutionary timescales. However, many lesser-studied groups, particularly those in the tropics, lack a densely sampled phylogeny which hinders robust inference of their evolutionary and biogeographic history. This study investigates the global diversification and systematics of the putative pantropical moth genus *Parasa* Moore (Lepidoptera: Limacodidae), with the aim of assessing the relative importance of dispersal and vicariance in shaping its distribution. Medium-coverage whole genome sequencing of specimens predominantly from museum collections were used to generate a globally sampled time-calibrated phylogeny of *Parasa* and associated genera (the *Parasa*-complex). Ancestral range estimation analyses were employed to infer geographical origins and possible dispersal times between bioregions. The *Parasa*-complex originated in Africa in the late Oligocene (∼24 Ma) and, through a series of long-distance dispersal events during the early-mid Miocene, expanded into Asia (∼23 Ma) and the Americas (∼21 Ma). Across all regions, dispersal was the dominant process shaping present-day distributions, with a limited role of vicariance in some subregions. Phylogenetic analyses further demonstrated that *Parasa* is not a monophyletic genus, with multiple independent lineages contributing to its apparent pantropical distribution. These findings highlight a central role of long-distance dispersal in generating certain global distributions. The results support a dynamic model of range evolution involving rapid Miocene dispersal and subsequent regional diversification. In addition, the non-monophyly of *Parasa* requires substantial taxonomic revision, underscoring the importance of robust phylogenetic frameworks for interpreting global biodiversity patterns.

## Introduction

Species vary widely in their distributional extent, from narrow range endemics to globally distributed taxa, and understanding this variation is a central theme of biogeography. Contemporary biogeographic theory recognises that both vicariance and dispersal jointly shape species distributions, rather than acting as competing, mutually exclusive processes (Sanmartín, 2012; Lomolino *et al*., 2016; Bourguignon *et al*., 2018). Traditionally, vicariance has been invoked as the dominant explanation for taxa with broad, disjunct distributions spanning multiple continents, particularly where lineage divergence times coincide with major geological events such as the breakup of Gondwana (Rosen, 1978; Pirie *et al*., 2015). However, an increasing number of studies are challenging vicariance-heavy interpretations of global distribution patterns, instead highlighting a substantial role of long-distance dispersal (Harris *et al*., 2018; Conceição Oliveira *et al*., 2021) particularly amongst taxa with strong dispersal capabilities (Torres-Cambas *et al*., 2019). As a result, there is a growing need for studies of widely distributed taxa to disentangle the relative contributions of these processes across spatial and temporal scales.

Taxonomic groups exhibiting broad but disjunct distributions offer model systems to test drivers of distributional patterns through Earth’s geological history (Leong *et al*., 2025). There are numerous examples of widespread or pantropical genera which exist in disjunct tropical regions across the globe, including plants (Rabeau *et al*., 2017; Jiao *et al*., 2025; Ang *et al*., 2025; Nge *et al*., 2026), insects (Economo *et al*., 2015; Fric *et al*., 2019) and mammals (Christiansen, 2008). Studies of these systems have demonstrated that pantropical distributions can be explained by vicariance (Kim & Farrell, 2015; Toussaint *et al*., 2017), long-distance dispersal (Nie *et al*., 2013; Fric *et al*., 2019; Conceição Oliveira *et al*., 2021; de Mestier *et al*., 2022; Smith *et al*., 2026; Domenech *et al*., 2026) or a combination of both (Jurado-Rivera *et al*., 2017; Ye *et al*., 2017; Toussaint & Short, 2018; Takayama *et al*., 2021). Beyond biogeographic inference, such broadly distributed taxa can also illuminate how geological events and ecological processes shape evolution. For example, the closure of the Tethys Ocean led to lineage diversification in a large and ecologically important pantropical mangrove genus (Takayama *et al*., 2021), whilst shared selective environmental pressures on different continents drive intraspecific trait clines in grasses (Møller *et al*., 2026). Together, such research builds our understanding of the processes shaping global biodiversity patterns.

Robust interpretations of dispersal, vicariance and trait evolution depend on densely sampled, time-calibrated phylogenies wherein taxonomic boundaries are clearly established. Prior to the advent of molecular sequencing, taxonomic classification relied on morphological traits. However, parallel adaptive evolution often produces phenotypic similarity and homoplastic characters in unrelated taxa (Losos, 2011; Stayton, 2015). Molecular phylogenetics has demonstrated such phenotypic convergence in globally distributed taxonomic groups, which were subsequently found to be non-monophyletic (e.g., Helbig *et al*., 2005; Haring *et al*., 2007; Alström *et al*., 2011; Cai *et al*., 2019; Ringelberg *et al*., 2022; Larson *et al*., 2023; Cordes *et al*., 2024; Leong *et al*., 2025). Such taxonomic inaccuracies can mislead inferences about diversification dynamics, ancestral trait evolution and biogeographical history. However, phenotypic information can highlight cases of repeated, independent evolution where similar traits are favoured across geographical realms.

Lepidoptera represent one of the most diverse and ecologically significant insect groups, possessing strong dispersal abilities and found on all continents except Antarctica. Due to these features, they are particularly informative for testing mechanisms of global distributions. Genera (or even species) of Lepidoptera that are extremely widespread tend to have migratory tendencies, exist in temperate regions and/or have recently diverged (Palahí *et al*., 2025), but those that are truly pantropical with supported monophyly are rare (Taberer, 2024). Furthermore, studies of diversification and biodiversity of many moth genera remain scarce on an intercontinental scale (Murillo-Ramos *et al*., 2026). The widespread moth genus *Parasa* Moore (Lepidoptera: Zygaenoidea: Limacodidae) is found throughout the global tropics in south and central America, Africa, Madagascar and southern and eastern Asia, as well as some sub-tropical/temperate zones in the Nearctic and parts of the Palearctic and western Asia. Members of the genus typically share a distinctive phenotype characterised by a green and brown banded forewing pattern (Fig. 1). Additional genera have been associated with *Parasa* due to morphological similarities, typically in the green forewing colouration and/or larval morphology (Solovyev, 2014; Epstein *et al*., 2025). This group is known as the *Parasa*-complex, of which there are around 240 species (Lin *et al*., 2019).

**Figure 1.**
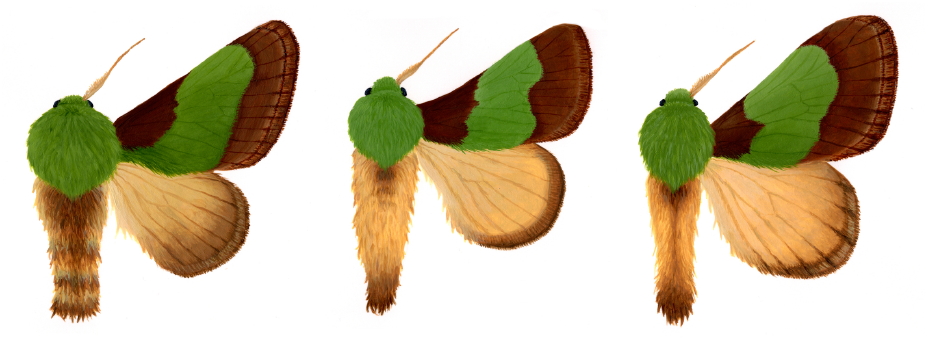
Phenotype of species within the *Parasa*-complex, all originally described in the genus *Parasa*, from disjunct global regions. Left: *Parasa chloris* (North America); Middle: *Parasa ananii* (West Africa); Right: *Aergina hilaris* (South Asia).

Although *Parasa* has long been described as pantropical, multiple studies have suggested that the genus as currently defined is not monophyletic, even within continents (Solovyev, 2014; Zaspel *et al*., 2016; Lin *et al*. 2019; Epstein *et al*., 2025). Despite such prior attention, no work has attempted to infer the evolution and biogeographic history of this group on a global scale. The *Parasa*-complex lacks a densely sampled phylogeny, with particular under-representation from the Afrotropics (e.g., Zaspel *et al*., 2016; Lin *et al*. 2019). Recent work by Epstein *et al*. (2025) represents the best-sampled phylogeny to date, although this work was focused on the higher classification of the family Limacodidae.

Here, medium coverage whole genome sequencing (WGS) and time-calibrated phylogenetic reconstruction with global sampling of *Parasa* and its associated genera is used to i) test the monophyly of this putative pantropical genus; infer the likely geographic origin of the group; evaluate the relative roles of dispersal and vicariance in shaping its global distribution; and iv) comment on the taxonomic classification of *Parasa*.

## Methods

### Taxon sampling

Three broad geographical bioregions are considered: Americas, Afrotropics and Asia. Alongside species currently classified within *Parasa* itself, various members of the *Parasa-*complex were included to determine their relation to the genus. The Asian *Parasa*-complex (encompassing Indomalaya and the Palearctic) was revised in Solovyev (2014) to also contain *Thespea* Solovyev, *Circeida* Solovyev, *Nephelimorpha* Solovyev, *Aergina* Solovyev, *Thronia* Solovyev, *Canon* Solovyev, *Melinaria* Solovyev, *Soteira* Solovyev, *Laphridia* Solovyev, *Polyphena* Solovyev and *Caiella* Solovyev. For the *Parasa* of the Americas, current knowledge is based on Epstein (1996) and Epstein & Corrales (2004), which list the following genera as members of the genus-complex: *Euclea* Hübner, *Acharia* Hübner, *Adoneta* Clemens, *Monoleuca* Grote & Robinson, *Talima* Walker, *Paraclea* Dyar and *Zaparasa* Dyar.

There is no recent revision of Afrotropical members of the genus, although many publications have considered *Latoia* Guérin-Méneville as closely associated to or synonymous with *Parasa* (e.g., Inoue, 1970; Epstein & Corrales, 2004; Solovyev, 2014). Through careful study of specimens held at ANHRT (African Natural History Research Trust, Leominster, UK), a rich source of African Lepidoptera, as well as the literature, the following genera were identified as Afrotropical members of the *Parasa*-complex: *Latoia, Ximacodes* Viette, *Parnia* Mabille, *Hilipoda* Karsch, *Stroter* Karsch, *Delorhachis* Karsch and *Letois* Felder & Felder. Outgroups were selected as members from three other Limacodidae subfamilies: Chrysopolominae, Crothaeminae and Dalcerinae. Identification and current taxonomy were based on the most recent literature available for the group (e.g., Solovyev, 2014 for Asian *Parasa*) as well as museum curation.

In total, 63 species of the *Parasa*-complex and four outgroup species were included. All genera of the *Parasa*-complex were represented in this study except for *Zaparasa* and *Canon*, due to a lack of recent samples in museums and/or sequencing success. For full details of species and information on specimens included in this study, see Table S1.

Specimens were located and loaned from several institutions: ANHRT, MNHN (Muséum National d’Histoire Naturelle, Paris, France) USNM (Smithsonian Institution, Washington D.C., USA) and ZSM (Zoologische Staatssammlung München, Munich, Germany). In most cases two samples of each species were loaned for DNA extraction to ensure sufficient DNA amount and quality. Specimens were selected on the basis of age, as younger, more recently collected material generally performs much better than older samples for DNA extraction and sequencing (Miller *et al*., 2013; Blaimer *et al*., 2016; Mayer *et al*. 2021). One specimen of *Parasa dusii* Solovyev & Saldaitis from the Arabian region was obtained from the private research collection of Stefano Dusi (Verona, Italy). In total, 62 museum specimens were sequenced for this study.

Five specimens included in this study were collected and immediately stored in >80% ethanol from opportunistic field sampling events. From the U.S.A., a single *Parasa chloris* (Herrich-Schäffer) specimen was raised from ova in George Washington University from a wild-caught female from Perry County, Ohio, and a single *Parasa indetermina* (Griffith & Pidgeon) specimen was collected from Cape May, New Jersey. From the Republic of Congo, three specimens were included as part of collecting expeditions by ANHRT in Noubalé-Ndoki National Park (see Takano, 2024) for more details on the expeditions).

### DNA extraction, library preparation and sequencing

All museum specimens and their labels were imaged. Most of the abdomen was removed, sterilising forceps and microscissors with >96% ethanol and waving over a flame in between each sample. The final segments of the abdomen were saved in labelled plastic micro vials for the later potential dissection of the genitalia, and the remaining segments of the abdominal tissue were taken forward for DNA extraction. Use of the abdominal tissue was no more destructive than regular practises of genital dissection of museum samples for taxonomic purposes (Strutzenberger *et al*., 2012; Mayer *et al*., 2021), enabling genitalia dissections for species identification and morphological data if required in the future, whilst maximising the amount of tissue used for DNA extraction and sequencing (Hebert *et al*., 2013).

The dried tissue was first rehydrated for approximately ten minutes in ∼150 µL ultrapure water. DNA was extracted using the Qiagen DNeasy Blood & Tissue kit (Qiagen, Hilden, Germany) and the standard DNA extraction protocol for tissue was followed (incubating at 56°C for ∼16 hours overnight) with the following modifications: 4 µL RNase (100 mg/ml) was added after the digestion step to destroy any RNA present in the samples, and the final elution was concentrated by two successive steps of adding 100 µL of Buffer AE to the spin column. DNA concentration was quantified with a Qubit 2.0, and fragment lengths were inspected with a 1.5 % agarose gel electrophoresed at 80V for 45 minutes.

The extracts were sent to Novogene UK for library preparation using the Novogene NGS DNA Library Prep Set. Genomic DNA was randomly sheared into short fragments, however, if most of the fragments were between 200 and 400 bp, this shearing step was skipped. The fragments were end repaired, A-tailed and further ligated with Illumina adapters. These were then PCR amplified, size selected and purified. The library was checked with Qubit and real-time PCR for quantification and with a bioanalyzer for size distribution detection. Quantified libraries were pooled and sequenced as 150 bp paired-end reads to an intended depth of 10X coverage on the Illumina Novaseq X Plus platform (Illumina, San Diego).

### Data quality and filtering

Quality of raw reads were first assessed with FastQC (http://www.bioinformatics.babraham.ac.uk/projects/fastqc) and then trimmed to remove adapter content and base calls of low quality using fastp (Chen *et al*., 2018) with default settings and 10 bp trimmed from the start of each read. Due to there being no appropriate reference genome available, genomes were assembled *de novo* using SPAdes v4.2.0 (Bankevich *et al*., 2012) and the quality of the resulting assembly assessed with quast (Gurevich *et al*., 2013). *De novo* genome assemblies were processed with buscophy (https://gitlab.leibniz-lib.de/smartin/buscophy) with flag *-f*, which identified and extracted complete and fragmented Benchmarking Universal Single-Copy Orthologs (BUSCOs) (Manni *et al*., 2021) using a reference sample set of 5,286 Lepidoptera genes based on OrthoDB v.10 (Kriventseva *et al*., 2019) (dataset *lepidoptera_odb10*). This produced alignment files for each gene, with 5,081 identified in total. Number of bases and reads before and after trimming, details on contigs and scaffolds after *de novo* assembly and BUSCO gene recovery per sample can be found in Table S1 and Fig. S1.

OliInSeq (https://github.com/cmayer/OliInSeq) was used to remove outliers from the corresponding amino acid and nucleotide alignments. Any columns in the alignments that had more than 50% missing data were removed with trimAl (Capella-Gutiérrez *et al*., 2009). Alignments were then further filtered in Geneious Prime 2025.0.3 (http://www.geneious.com/) to remove those with less than 50% of samples represented, those with a GC content over 60% (see Bossert *et al*., 2017), those with a pairwise identity of over 98%, and those with a sequence length of less than 400 bp. This left 2,024 gene alignment files. Summary statistics for each alignment created using AMAS (Borowiec, 2016) are in Table S2.

### Phylogenetic analyses

Phylogenetic tree searches were completed using two methodologies. Firstly, the gene alignments were concatenated using AMAS, creating an alignment of 3,229,490 bp. The concatenated alignment and partition file were inputted into IQ-TREE v1.6.12 (Nguyen *et al*., 2015) to first search for the best model for each partition with *-m MFP*. Next, in the tree search stage in IQ-TREE v2.2.2.6 (Minh *et al*., 2020), the resulting scheme.nex file was inputted as the *-p* and *-m* flag. Branch support was calculated with 1000 ultrafast bootstrap replicates *(-bb 1000*) (Minh *et al*., 2013) and approximate likelihood-ratio test (*-alrt 1000*) (Anisimova & Gascuel, 2006). The bootstrap trees were optimised with *-bnni*, and tree searches were run 10 times independently to search for the best-scoring tree (*--runs 10*). This resulting tree is referred to as the IQ-TREE phylogeny herein.

In the second method, IQ-TREE v2.2.2.6 was used to generate individual trees from each gene alignment. Branch support was calculated with 1000 ultrafast bootstrap replicates and approximate likelihood-ratio test, and with five likelihood searches for each alignment (*--runs -5*) to produce the best-scoring tree. These gene trees were then combined with ASTRAL-IV (within package ASTER; Zhang *et al*., 2025) to form a consensus phylogeny. This is referred to as the ASTRAL phylogeny herein.

Concordance factors were tested due to the potential for gene tree discordance when utilising numerous loci, as each gene may produce a distinct tree topology that is different to the species tree (Lanfear & Hahn, 2024). ASTRAL was used to test for quartet concordance (qCF) with *-t 2*, and IQTREE was used to test for gene (gCF) and site (sCF) concordance with *--gcf* and *--scfl 100*, respectively. Results suggested that in general, gCF was higher than sCF across the branches and hence the ASTRAL phylogeny is displayed as the main result (Fig. 2). Concordance calculations can be found in Table S3. Despite better concordance for the ASTRAL phylogeny, the overall topology produced by both methods were almost identical (discussed below; see Fig. S2).

**Figure 2.**
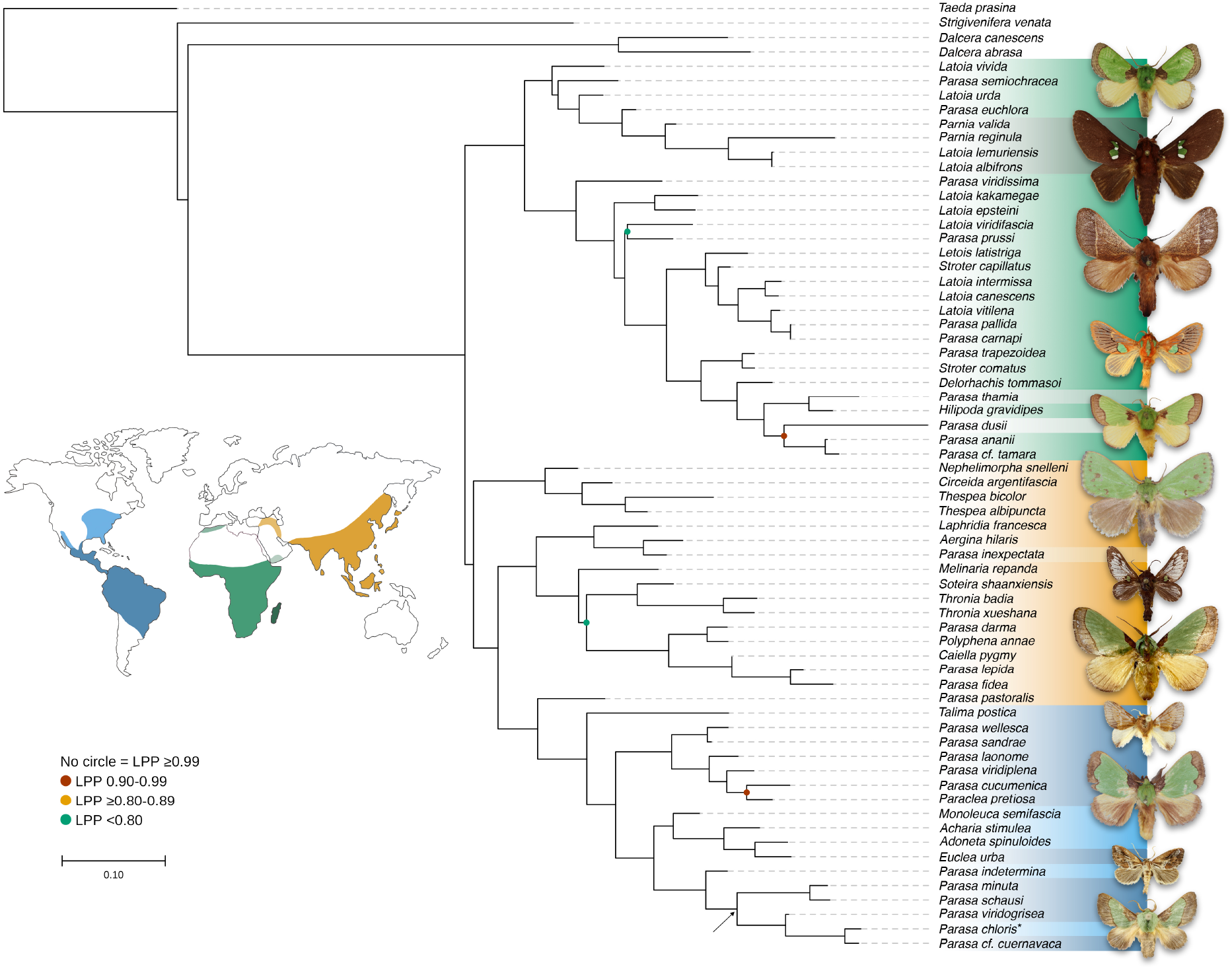
ASTRAL phylogeny of the global *Parasa*-complex (excluding *Ximacodes*). Constructed with ASTRAL-IV (within package ASTER; Zhang *et al*., 2025) by combining 2,024 individual BUSCO Lepidopteran gene trees made in IQ-TREE v2.2.2.6 (Minh *et al*., 2020). Branch local posterior probability (LPP) is given in the key. Biogeographic regions represented with map colours. Asterisk denotes type species of *Parasa, Parasa chloris*. Arrow points to clade putatively considered *Parasa sensu stricto* (see Discussion: Taxonomy of *Parasa sensu lato*).

The sample for *Ximacodes affinis* (Mabille) had the lowest base pair recovery, with just 14 complete or fragmented BUSCO genes remaining after data filtering. The above analyses were therefore repeated but with *X. affinis* removed to test if branch support improved, resulting in four final phylogenies (see Fig. S2 for the phylogenies and Table S4 for concordance calculations). The ASTRAL phylogeny without *Ximacodes* is shown in Fig. 2.

### Divergence time estimations

There are relatively few Lepidopteran fossils compared to other insects (Labandeira & Sepkoski, 1993), and no well-identified Limacodidae (Sohn *et al*., 2012). For this reason, the tree utilised two secondary calibration points from the recent fossil-calibrated Zygaenoidea study by Mirić *et al*. (2024): one between two of the outgroups, and one at the root. Due to the size of the dataset, divergence times of the *Parasa*-complex were estimated with LSD2 v2.4.1 (https://github.com/tothuhien/lsd2), a rapid least-squares algorithm. For this, the IQ-TREE phylogeny was used as the constraint topology as LSD2 requires a tree with branch lengths in substitution units. The calibration point was used on the outgroups *Dalcera* Herrich-Schäffer (Limacodidae: Dalcerinae) + *Strigivenifera* Hering (Limacodidae: Chrysopolominae) (72.5–55.7 Ma) based on the minimum and maximum ages obtained by Mirić *et al*. (2024). In addition, a constraint at the root of the phylogeny of 90.0– 81.0 Ma was given as the age of the superfamily Zygaenoidea, also based on Mirić *et al*. (2024). LSD2 was configured to preserve the input tree topology by setting the informative branch length threshold to zero *(-l* 0) and enforcing minimum branch lengths (*-u* e) to prevent potential collapse of short branches into polytomies. To calculate confidence intervals, 100 bootstrap trees were simulated with *-f* 100 (Fig. S3).

For comparison with the least squares approach, divergence times were also estimated using treePL, a penalised-likelihood approach (Smith & O’Meara, 2012). Methodology partly followed Kawahara *et al*. (2023) whereby the initial priming step was run 100 times on the starting tree to find the majority *opt, optad* and *optcvad* values (as with LSD2, the IQ-TREE phylogeny was used as the input tree). These were then inputted into the configuration file before running the cross-validation step 100 times to find the majority smoothing value. Cross-validation was performed with 10 replicates, 100,000 simulation iterations, and smoothing values decreasing from 10 to 0.0001. To obtain confidence intervals, 100 bootstrap trees were estimated in IQ-TREE using the IQ-TREE phylogeny as a topology constraint so the bootstrap trees were compatible with each other. The best smoothing value was then added as a variable to the final step, which involved running treePL once on each bootstrap replicate. A consensus tree file was generated with TreeAnnotator v.2.7.7 (Drummond & Rambaut, 2007), with 0% of burnin and mean node heights, following Maurin (2020) (Fig. S4).

### Biogeographic analyses

Species distributions were assigned as present/absent to the eight following biogeographical subregions, as defined from the three broad bioregions: Neotropic and Nearctic (Americas); West Palearctic, Africa, Madagascar and Arabia (Afrotropics); West Asia and East Asia-Indomalaya (Asia) (Table S1; see map within Fig. 2 and Fig. 3). In this case, West Palearctic refers to just northern Morocco and northern Algeria and is hence considered a subregion of the Afrotropics. Ancestral area estimations were performed in R v4.3.2 (R Core Team 2023) using the package *BioGeoBEARS* (Matzke, 2013), whereby a model testing approach was used to test the fit of three commonly used biogeographic inference models: Dispersal-Extinction-Cladogenesis (DEC; Ree & Smith, 2008), BAYAREALIKE (Landis *et al*., 2013) and DIVALIKE (Ronquist, 1997). These models were implemented with and without considering the jump parameter for founder events (+*j*; Matzke, 2014). The dated phylogeny built with LSD2 was used as input, and the maximum number of areas a taxon could occupy was two.

**Figure 3.**
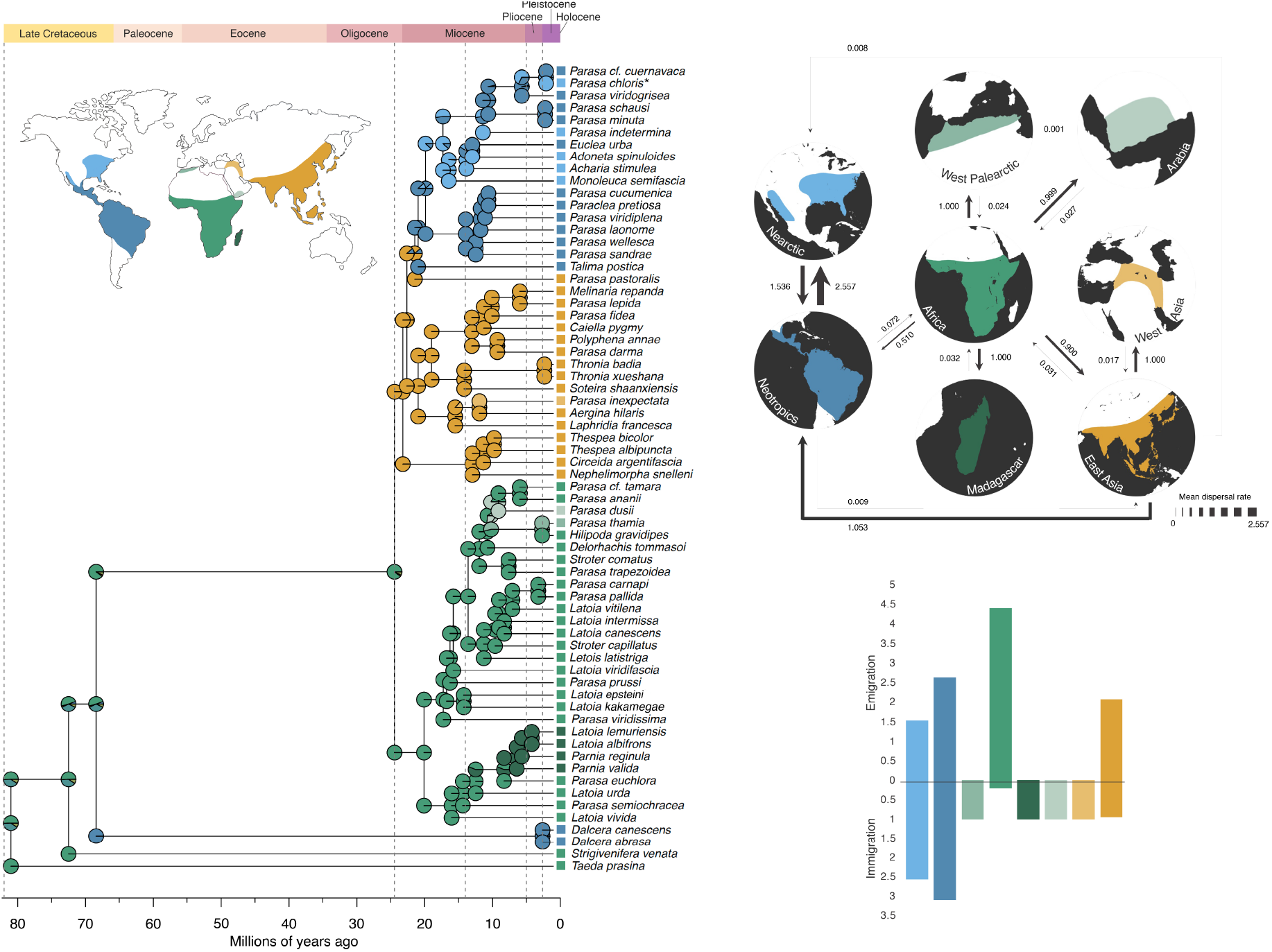
Left: Ancestral range reconstruction of the global *Parasa*-complex based on the time calibrated DEC+*j* model as implemented in *BioGeoBEARS* (Matzke, 2013). Alternative inferences of ancestral ranges are provided in Figure S6. The chronogram used as input was created using LSD2 v2.4.1 (https://github.com/tothuhien/lsd2) (see Methods). Coloured boxes at each terminal name represent the eight biogeographical subregions as illustrated in the upper-left map. Top right: Relative mean dispersal rates between bioregions based on 1000 simulations under the DEC+*j* model; arrows between regions indicate the direction of each dispersal event, and arrow thickness is scaled to the relative mean dispersal rate. Bottom right: Mean immigration and emigration frequencies of each bioregion based on 1000 simulations under the DEC+*j* model.

These ancestral area estimations were run under two conditions: (1) an unconstrained analysis with no time stratification nor different dispersal scalars between regions; (2) a time-stratified analysis with dispersal rate scalars to account for changes in geography over time. For (2) the time periods were: 0–2.6 Ma (present to start of Pleistocene), 2.6–5 Ma (Pliocene), 5–14 Ma (mid-to late-Miocene), 14–24.4 Ma (mid-Miocene to late-Oligocene, and crown age of *Parasa*-complex), and 24.4 Ma–82 Ma (late-Oligocene to late-Cretaceous; maximum phylogeny date). Dispersal probabilities among geographic areas were estimated by observing distances between landmasses in geological reconstructions using the R package *rgplates* v0.6.1 (Kocsis *et al*., 2024) under the plate rotation model of Müller *et al*. (2019) (Fig. S5). Dispersal rate matrices between areas (Table S5) and results of model selection (evaluated with log-likelihood values and AICc scores) are shown in Table S6 and Fig. S6.

The number and type of biogeographic event was estimated using biogeographical stochastic mapping (BSM) in *BioGeoBEARS*. 1000 simulations were run under the DEC+*j* model, and the relative mean dispersal rates were calculated to obtain event frequencies of immigration and emigration (Fig. 3; Table S7).

## Results

### Phylogenetic trees and topology

The recovered phylogenies revealed that the genus *Parasa* was polyphyletic with respect to other genera in the *Parasa*-complex across its global distribution (Fig. 2, Fig. S2). In the ASTRAL phylogeny without *Ximacodes*, branch support was generally very high, with most branches receiving a local posterior probability (LPP) of ≥0.99 (Fig. 2). Two branches had LPP between 0.90−0.99, the first containing three *Parasa* species found in the Afrotropical realm and the second containing a *Parasa* species + *Paraclea* found in the Neotropics. There were two branches with low support (LPP <0.80), one containing a species of *Latoia* and a species of *Parasa* in Africa (LPP=0.32), and the second containing *Soteira* + *Thronia* + *Polyphena* + *Caiella* + three members of *Parasa*, all from Asia (LPP=0.41). In the corresponding IQ-TREE phylogeny (i.e. without *Ximacodes*; Fig. S1), branch support was extremely high with both ultrafast bootstrapping (uBS) and approximate likelihood ratio test (SH-aLRT) of ≥99 for all nodes but one where uBS ≥80−99 and SH-aLRT ≥90.0−99.0.

This node contained some members of *Parasa, Latoia* and several other genera, all found in the Afrotropics.

When including *Ximacodes* (Fig. S2), several more branches in the ASTRAL phylogeny received LPP <0.99, indicating that its presence introduced some uncertainty. In the corresponding IQ-TREE phylogeny, branch support values were unchanged from the phylogeny excluding *Ximacodes* (the branch containing *Ximacodes* received support of uBS=100 and SH-aLRT=100.0). In both analyses, *Ximacodes* was not placed with other taxa from Madagascar, but instead was sister to a clade containing nine *Parasa* taxa + six *Latoia* taxa + *Letois* + *Stroter* + *Delorhachis* + *Hilipoda* (all Afrotropical).

The topology of all four phylogenies was extremely similar, with only the position of *Melinaria* differing (Fig. S2). The topology generally shows a split between Afrotropical taxa and Asian + American taxa. *Melinaria* was recovered as sister to a clade containing *Soteira* + *Thronia* + *Polyphena* + *Caiella* + three *Parasa* taxa in the ASTRAL phylogenies, but as sister to just *Parasa lepida* in the IQ-TREE phylogenies (all Asian taxa). Any other conflicts, as shown in Fig. S2, were pairwise branch rotations which did not affect tree topology.

### Divergence time estimates and biogeography

According to time calibration with LSD2 (Fig. S3), the clade containing members of the *Parasa*-complex originated around 24.4 Ma (28.4–19.4 Ma) (Late Oligocene/Early Miocene), after which it split into two clades (Afrotropical and Asia + America) that simultaneously and rapidly diversified. A similar timing of origin of the complex was found in the treePL calibrated phylogeny of 24.3 Ma (24.2–24.5 Ma) (Fig. S4), although the confidence intervals associated with the timing estimate were extremely narrow.

The best supported model of ancestral range reconstruction with *BioGeoBEARS* was the time-stratified DEC+*j* model, which received the highest log-likelihood value and the lowest AICc score (Fig. 3; Table S6). This model, alongside the BSM results, supported Africa as the most likely inferred ancestral area occupied by the most recent common ancestor (MRCA) of the *Parasa*-complex. The model then suggests dispersal from Africa into East Asia-Indomalaya around 23 Ma (26.9– 18.5 Ma). From here, dispersal occurred into the Americas almost 21 Ma (25.0–16.6 Ma), with mean dispersal rates from East Asia-Indomalaya higher to the Neotropics (1.053) than the Nearctic (0.008).

In all time stratified analyses, model fit was improved compared to each counterpart unconstrained model. Without inclusion of the *j* parameter for founder effects, the time-stratified DIVALIKE model had the best fit, but inferred a less certain ancestral range from either Africa or Africa + Asia (Table S6). When the *j* parameter was included, all ancestral range estimation unambiguously supported an African origin for the *Parasa*-complex (Fig. S6).

Within the Afrotropics, the African lineage dispersed to Madagascar, Arabia, and the West Palearctic. Taxa from Madagascar were recovered in a single clade resulting from a colonisation around 12 Ma (15.7–10.2 Ma) and subsequent diversification 6.4 Ma (8.5–5.0 Ma). Dispersal to Arabia by the lineage represented by the single extant taxon *P. dusii* occurred around 10 Ma (12.5– 7.8 Ma), with some indication of a reverse colonisation to Africa in the late Miocene, 6.0 Ma (7.9–4.2 Ma). Finally, *Parasa thamia* Rungs, the only extant *Parasa* species found in the West Palearctic, shared a less certain ancestral range with its sister taxon *Hilipoda gravidipes* Karsch, being Africa and the West Palearctic with a split occurring 2.7 Ma (3.6–1.9 Ma).

## Discussion

The global distribution of the *Parasa*-complex is predominantly explained by long-distance dispersal, with limited evidence for vicariance. Time-calibrated phylogenetic and biogeographic reconstructions indicate an African origin in the late Oligocene, followed by rapid Miocene range expansion into Asia (∼23 Ma) and the Americas (∼21 Ma). The timing of divergence broadly coincided with ecological opportunities and drivers that could have facilitated dispersal. In addition, the results demonstrate that *Parasa* is not a monophyletic genus, indicating that its apparent pantropical distribution does not reflect a single, cohesive evolutionary lineage.

### Biogeography and divergence times

The phylogenetic and biogeographic analyses support an African origin for the *Parasa*-complex around 24.4 Ma. Africa exhibited the highest emigration and lowest immigration rates of all, suggesting it functioned primarily as a source region. This pattern is consistent with “Out-of-Africa” scenarios proposed for several other widespread Lepidopteran lineages (Kodandaramaiah & Wahlberg, 2007; Aduse-Poku *et al*., 2009; Sahoo *et al*., 2018). Diversification within the *Parasa*-complex appeared to increase near-simultaneously across bioregions during the Miocene, like the results of Nge *et al*. (2026) on a pantropical genus of Annonaceae, coinciding with the Mid-Miocene Climatic Optimum (a period of global climatic warming) which likely facilitated range expansion and lineage diversification (Aduse-Poku *et al*., 2022).

Dispersal from Africa to Madagascar occurred around 12 Ma (15.7–10.2 Ma), followed by *in situ* diversification on the island. This timing greatly post-dates Madagascar’s separation from mainland Africa (∼125 Ma; Rabinowitz & Woods, 2006) consistent with the notion that most colonisation events to Madagascar occurred after assuming its current position in the Early Cretaceous (Krüger, 2007; Rabinowitz & Woods, 2006). Ancestral range reconstruction suggests that the most recent common ancestor of the Madagascan taxa and their closest extant mainland relative, *Parasa euchlora* Karsch, may have been distributed in Madagascar. Although a back-colonisation of mainland Africa by the *P. euchlora* lineage is possible, based on its strictly West African distribution, this may be an artefact of taxon sampling. Examples of such back-colonisations are scarce in the literature but have been noted in *Leptotes* Scudder butterflies (Fric *et al*., 2019) as well as some plant groups (Bauret *et al*., 2017; Skema *et al*., 2023; Blanco-Gavaldà *et al*., 2025), reflecting a dynamic biogeographic history between island and mainland.

Dispersal into East Asia-Indomalaya occurred around 23 Ma and appears consistent with long-distance dispersal from Africa rather than stepwise expansion via the Arabian Peninsula or West Asia. However, given the extremely low extant diversity of the *Parasa*-complex in Arabia and West Asia (each represented by a single species, both of which were sampled here), it remains possible that these regions once played a more significant role as transient dispersal corridors, with subsequent extinction obscuring this signal. The timing of this dispersal overlaps broadly with the formation of the Gomphotherium land bridge following the collision of the African and Eurasian plates (∼20 Ma; Straume *et al*., 2025), which has been proposed as an important route for faunal exchange (Rage & Gheerbrant, 2020), although its role in the dispersal of the *Parasa*-complex remains uncertain.

Later exchanges between Africa and the Arabian Peninsula occurred during the late Miocene (∼10 Ma), a period of increased humidity intervals in Arabia and the presence of grasslands, rivers and lakes that may have facilitated dispersal (Markowska *et al*., 2025). The only extant representative of the *Parasa*-complex in Arabia, *P. dusii*, diverged from its African MRCA during this period and is now restricted to isolated montane habitats in southern Arabia, where subtropical rainforest refugia persist (Solovyev & Saldaitis, 2010). Similarly, *P. thamia* in the western Palaearctic is the only extant species of *Parasa* to be found in this region, splitting from its MRCA almost 3 Ma. This divergence from taxa found in West Africa’s Guinean forests may have been caused by the expansion of the Sahara Desert (Zhang *et al*., 2014; Armstrong *et al*., 2023), indicating that the disruption of forested corridors may have acted as vicariant barriers for these groups.

Dispersal from East Asia into the Americas is estimated to have occurred around 21 Ma, potentially via the Beringian land bridge, which intermittently connected these regions throughout the Miocene and Pliocene and has been widely implicated in intercontinental exchanges (e.g., Wen *et al*., 2016). However, dispersal was much higher from East Asia to the Neotropics as opposed to the Nearctic, which may be a consequence of incomplete taxon sampling or an artefact of founder-effect speciation as implemented by the +*j* parameter in *BioGeoBEARS*. The highest mean immigration rates overall were inferred from the Neotropics to the Nearctic, although movement in the opposite direction was also relatively high compared to other regions. Ancestral range reconstructions did not recover strong phylogenetic structuring between Nearctic and Neotropical taxa, suggesting possible repeated bidirectional exchange between temperate and tropical regions from the Miocene to the present. This pattern is consistent with evidence for a prolonged and complex process of the Great American Biotic Interchange beginning around 23 Ma (Bacon *et al*., 2015; Freitas-Oliveira *et al*., 2025), with symmetrical biotic exchanges between these two regions occurring until approximately 6 Ma (Bacon *et al*., 2015).

### Phylogenetic relationships

Results from this work based on WGS and a globally sampled phylogeny of the *Parasa*-complex reveal that *Parasa* is not monophyletic and thus cannot be considered a single widespread pantropical genus. Even within each of the three bioregions (Americas, Afrotropics and Asia), the genus recovered as polyphyletic, indicating the need for large-scale taxonomic revision. This result is consistent with previous work (Solovyev, 2014; Zaspel *et al*., 2016; Lin *et al*. 2019), and the placement of taxa is broadly similar to that of the *Parasa*-complex clade as defined in the Limacodid phylogeny by Epstein *et al*. (2025) based on seven molecular markers and 122 morphological characters. As with *Parasa*, the Afrotropical genera *Latoia* (n=9), *Stroter* (n=2) and *Parnia* (n=2) were also not recovered as monophyletic, suggesting that members of the *Parasa*-complex from this bioregion require systematic attention. The only other genera represented by more than one taxon, *Thronia* and *Thespea* (both Asian), were monophyletic, supporting the generic concepts of Solovyev (2014). For these reasons, the remainder of the discussion refers to this genus complex as *Parasa sensu lato* (*s*.*l*.).

It remains unclear whether the repeated occurrence of the green and brown banded forewing phenotype in *Parasa s*.*l*. arose convergently in multiple lineages or descended from a common ancestor with subsequent losses in intermediate genera. Resolving this question will require a more comprehensive phylogenetic framework, incorporating dense species-level sampling from all well-defined genera within the clade. In addition to the genera included in this study, Epstein *et al*. (2025) identified several others - *Latoiola* Hering, *Parapluda* Aurivillius, *Coenobasis* Felder & Felder, *Limacolasia* Hering and *Barisania* Holloway - as belonging to this broader clade. Until genus boundaries are more clearly defined, the evolutionary origins of this phenotype remain uncertain. It can be speculated that this phenotype may be ecologically advantageous due to its repeated appearance across several tropical regions. Green wing markings are relatively uncommon among moths, yet green colouration is seen in many other arthropods for camouflage in highly vegetated environments (Egorkin *et al*., 2025), like the tropics.

### Taxonomy of *Parasa sensu lato*

This study supports previous research in finding *Parasa* not monophyletic. Due to this, the taxonomy of this group of Limacodids remains unstable and requires major revision. As there are numerous well-defined, morphologically distinct genera recovered within the *Parasa*-complex clade (e.g., *Delorhachis*, as revised by Taberer *et al*., 2023), it would not be appropriate to synonymise all taxa in the present phylogeny into one genus. A more conservative solution would be to define the genus more strictly around the clade containing the type species and establish several new genera for the Asian and Afrotropical members of the genus complex.

Based on the results from this study and to stabilise global nomenclature, the first author is currently undertaking a large taxonomic revision of the Afrotropical *Parasa* with consideration of all taxa in the genus and associated genera within the region (>100 taxa). Within this work, several new genera will also be described from Asia. This builds on Solovyev (2014), who provisionally maintained numerous taxa within the genus *Parasa* as well-supported monophyletic species-groups, following a revision of the Palaearctic and Indomalayan members based on extensive morphological and limited genetic data. Whilst taxon sampling was not complete, these species-groups delimitations were recovered with the taxa included here, in Lin *et al*. (2019) and Epstein *et al*. (2025), and hence they likely represent good genera.

The taxonomy of American Limacodidae requires revision with a strict definition of *Parasa* based on the Nearctic type species *P. chloris*. Epstein *et al*. (2025) suggests that all genera from the Americas in this complex (including *Talima, Monoleuca, Acharia, Adoneta* and *Euclea*) could be synonymised with *Parasa*. However, there appears to be large morphological and genetic variation between these described genera which should be considered. One such synapomorphy to define the *Parasa sensu stricto* could be a paired cluster of setae as found in the male genitalia of *P. chloris* (illustrated in Taberer, 2025) and others in the same clade (marked with an arrow in Fig. 2). Such morphological feature is not observed in any *Parasa s*.*l*. in Asia or the Afrotropics (TRT pers. obs.), or even outside of this clade in the Americas, and thus makes a good candidate for genus delimitation alongside the genetic support provided here.

### Conclusion

Results here demonstrate how long-distance dispersal has acted as a main distributional driver of a global genus-complex, with rapid expansion across the tropics during the Miocene following an African origin. Whilst dispersal explains broad intercontinental patterns, subsequent climatic shifts leading to vicariant barriers have possibly functioned in structuring regional diversity. These findings reinforce a growing body of evidence that both dispersal and vicariance can play roles in shaping widespread tropical distributions.

The apparent pantropical distribution of *Parasa* does not reflect a single cohesive lineage, but instead comprises multiple, distantly related taxa belonging to distinct genera. This underscores the importance of densely sampled time-calibrated phylogenies for resolving the evolutionary history of widespread groups. The results also demonstrate the advantage of using museum specimens to sample lesser-studied groups from their entire tropical range, which would otherwise be challenging to obtain from nature. Finally, the work presents additional questions on the ecological advantages and biological pathways involved in the repeated evolution of the *Parasa*-like phenotype. As larger phylogenomic datasets become available on these lesser studied groups, our ability to test for shared ancestry or convergent adaptation to similar ecological pressures will greatly improve.

## Acknowledgements

The authors would first and foremost like to thank the museum curators who allowed access to the specimens under their care for this research and provided many interesting discussions about this project: Axel Hausmann (Zoologische Staatssammlung München, Munich, Germany), Scott Miller (Smithsonian Institution, Washington D.C., USA), Talitta Simoes (Smithsonian Institution, Washington D.C., USA) and Rodolph Rougerie (Muséum national d’Histoire naturelle, Paris, France). We also thank Aidas Saldatis (Institute of Ecology of Vilnius University, Vilnius, Lithuania) and Stefano Dusi (Verona, Italy) for allowing access to and organising transport for the *Parasa dusii* sample. In obtaining three field samples from the Republic of Congo, we are grateful to Hitoshi Takano (African Natural History Research Trust, Kingland, UK) for identifying and collecting the samples, and the Wildlife Conservation Society, Congo (WCSC) and their staff for authorising research and providing technical assistance. In particular, we thank Richard Malonga (Director, WCSC) and Morgane Cournarie (Program Coordinator, WCSC), Ben Evans (Park Director, PNNN), Vittoria Estienne (Director of Research and Biomonitoring, PNNN), Onesi Samba (Research Assistant, PNNN) and Yako Valentin (Administrative Manager, PNNN). The samples were collected under permit 051/MESRSIT/IRSEN/DG/DS and exported under permit 254/MESRSIT/IRSEN/DG/DS. We thank Joseph Goma-Tchimbakala (Directeur général, Institut National de Recherche en Sciences Exactes et Naturelles – IRSEN) for the seamless collaboration in issuing the necessary paperwork; to Jean Bosco Nganongo (Directeur, Direction de la Faune et des aires protégées) for issuing CITES permits; and Victor Mamonekene (Hydrobiologist, IRSEN and Marien Ngouabi University, Brazzaville) for his valuable assistance. Our thanks are extended to Luc Micheels (George Washington University, Washington D.C., USA) and John Lill (George Washington University, Washington D.C., USA) for their help in providing a sample of *Parasa chloris*, and for organising a field trip to Cape May, New Jersey to collect samples of *Parasa indetermina* (collected under Scientific Collecting Permit SC 2024139 - issued by the State of New Jersey Department of Environmental Protection Fish and Wildlife) for this project. Karina Lucas da Silva Brandao (Museum der Natur, Hamburg, Germany) is sincerely thanked for her assistance in access to specimens collected from Brazil. We also thank Alessandro Giusti (Natural History Museum, London, UK), Marc Epstein (California Department of Food & Agriculture, Sacramento, USA) and Alexey Solovyev (Ulyanovsk State Pedagogical University, Ulyanovsk, Russia) for their ongoing kindness and knowledge of Limacodidae which assisted with this work. DNA extraction and bioinformatic analysis could not have been possible without the initial assistance of Andrea Estandía (University of Oxford, Oxford, UK). Finally, the authors would like to acknowledge the use of the University of Oxford Advanced Research Computing (ARC) facility in carrying out this work (http://dx.doi.org/10.5281/zenodo.22558).

## Author contributions

TRT, TC and SMC conceived the idea for the project. TRT obtained specimens included in this study. Lab work was carried out by TRT with guidance from ME. TRT undertook bioinformatic analyses with guidance from ME and SM. TRT carried out further analyses and wrote the manuscript with input/editing from all other authors.

## Data availability statement

Sequence data generated for this work have been submitted to NCBI Sequence Read Archive under the accession number PRJNA1462647. Code used to analyse the read data is found at https://doi.org/10.5281/zenodo.20123356.

