## Supplementary figures and images for "Long-distance dispersal drives global tropical distributions in a widespread moth lineage (Lepidoptera: Limacodidae)"

### Figure S1

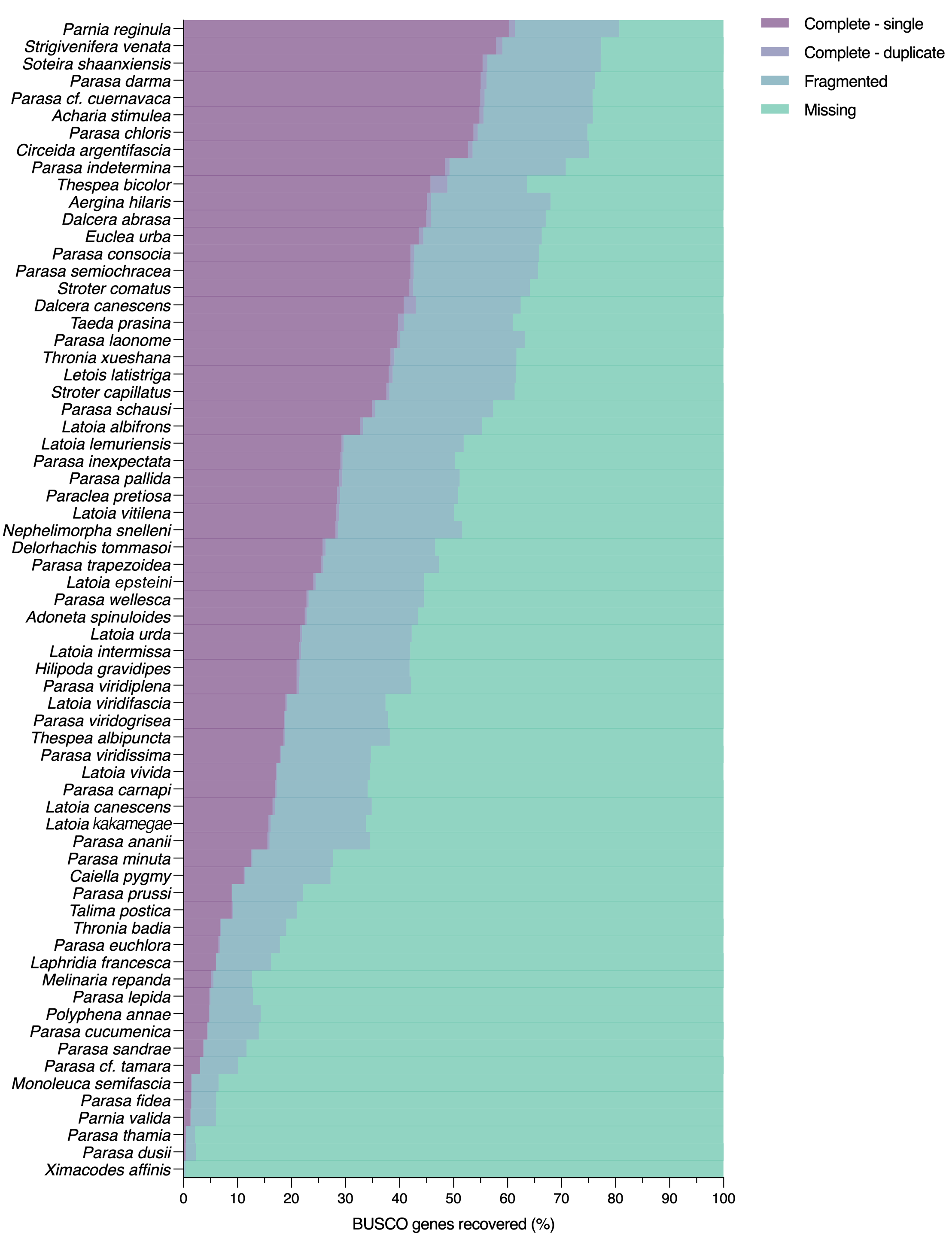

### Figure S2

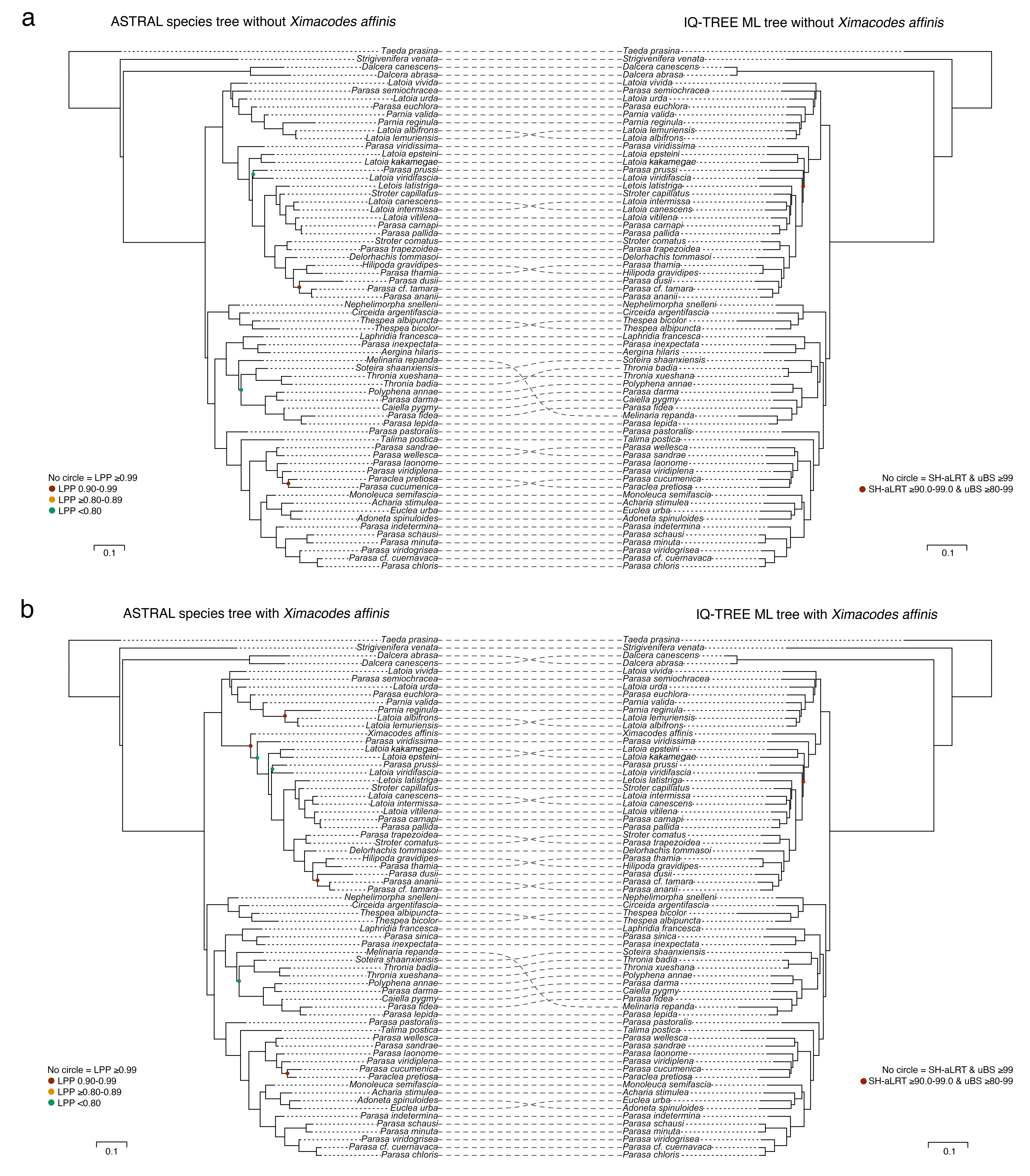

### Figure S3

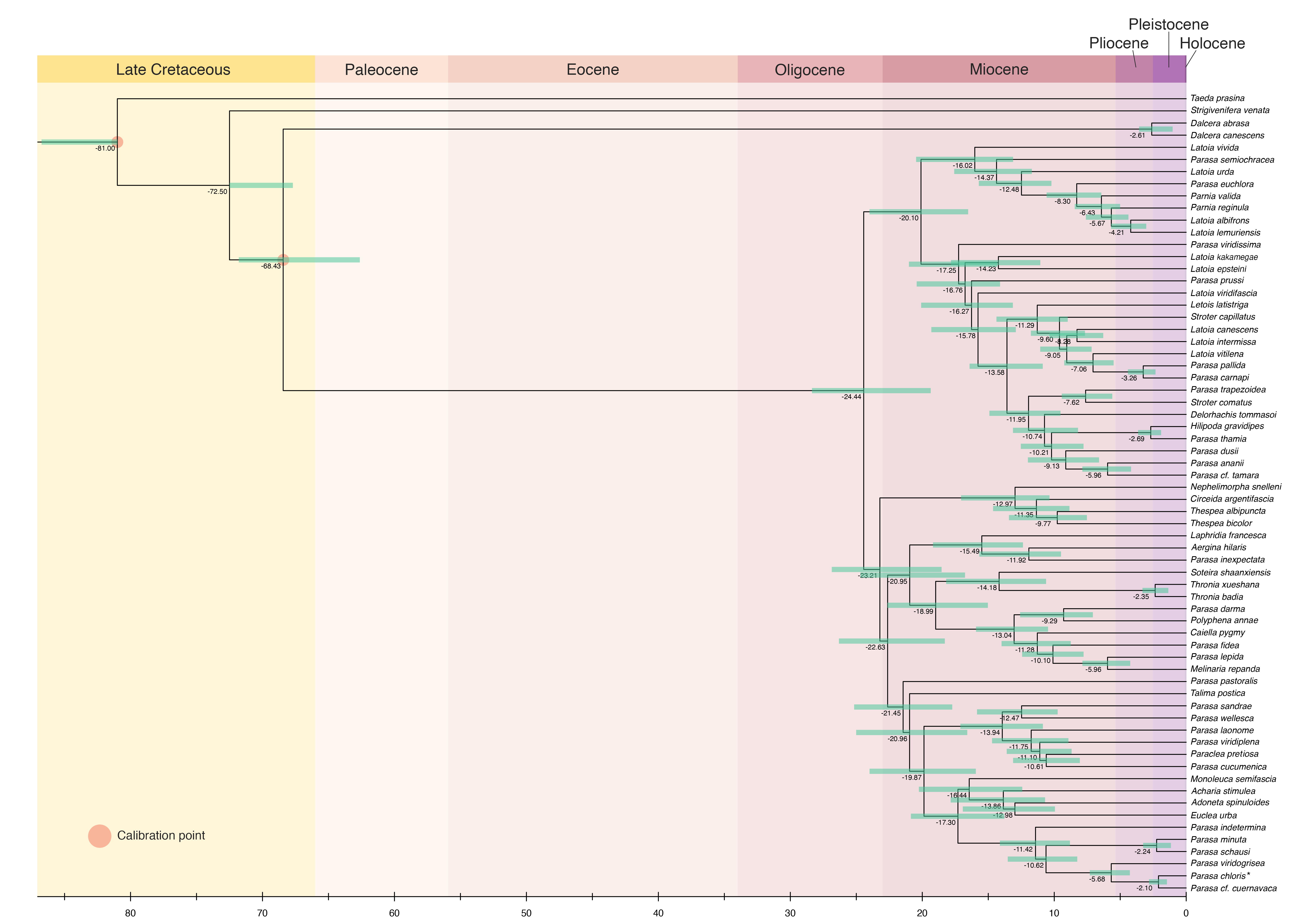

### Figure S4

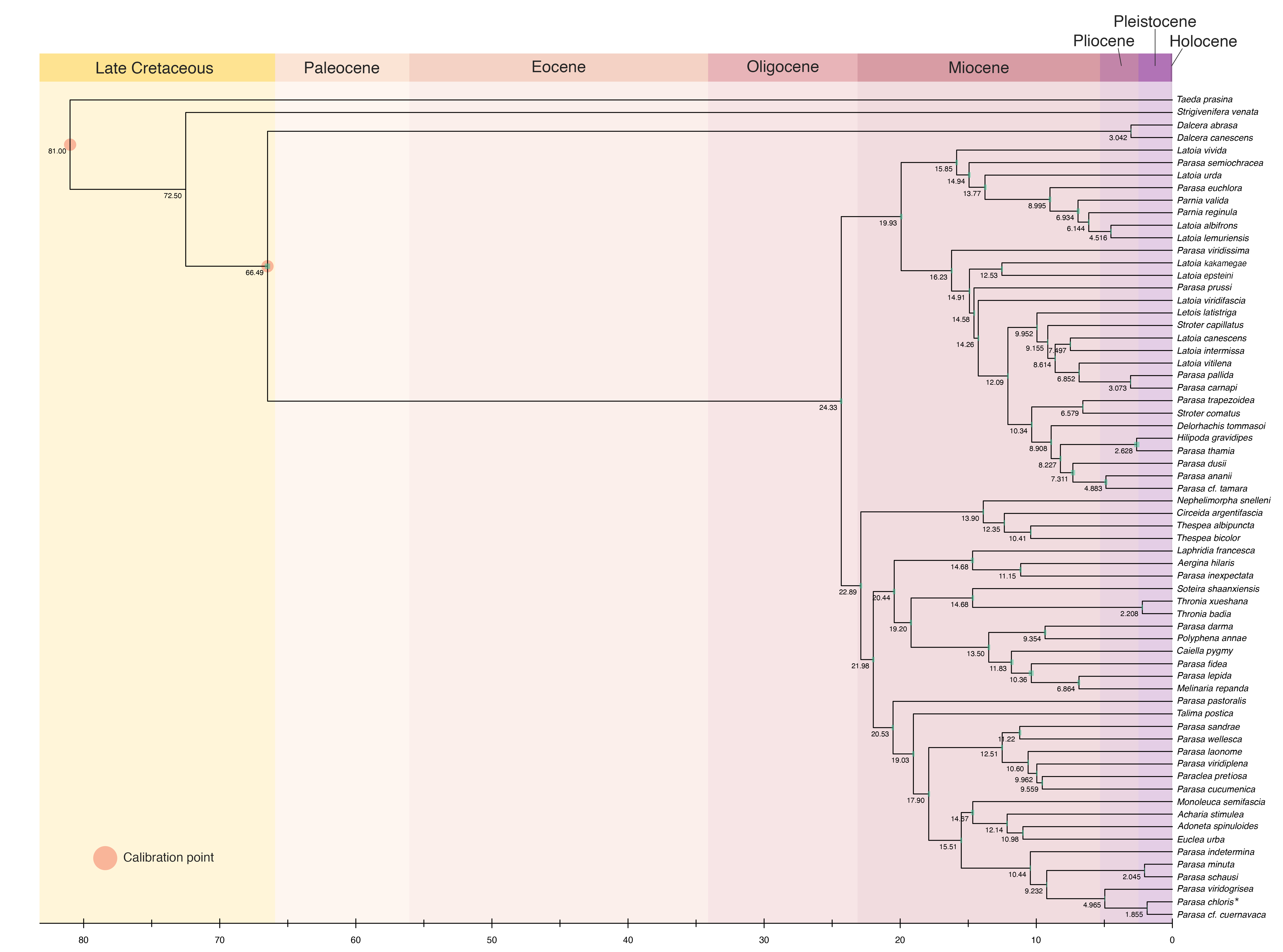

### Figure S5

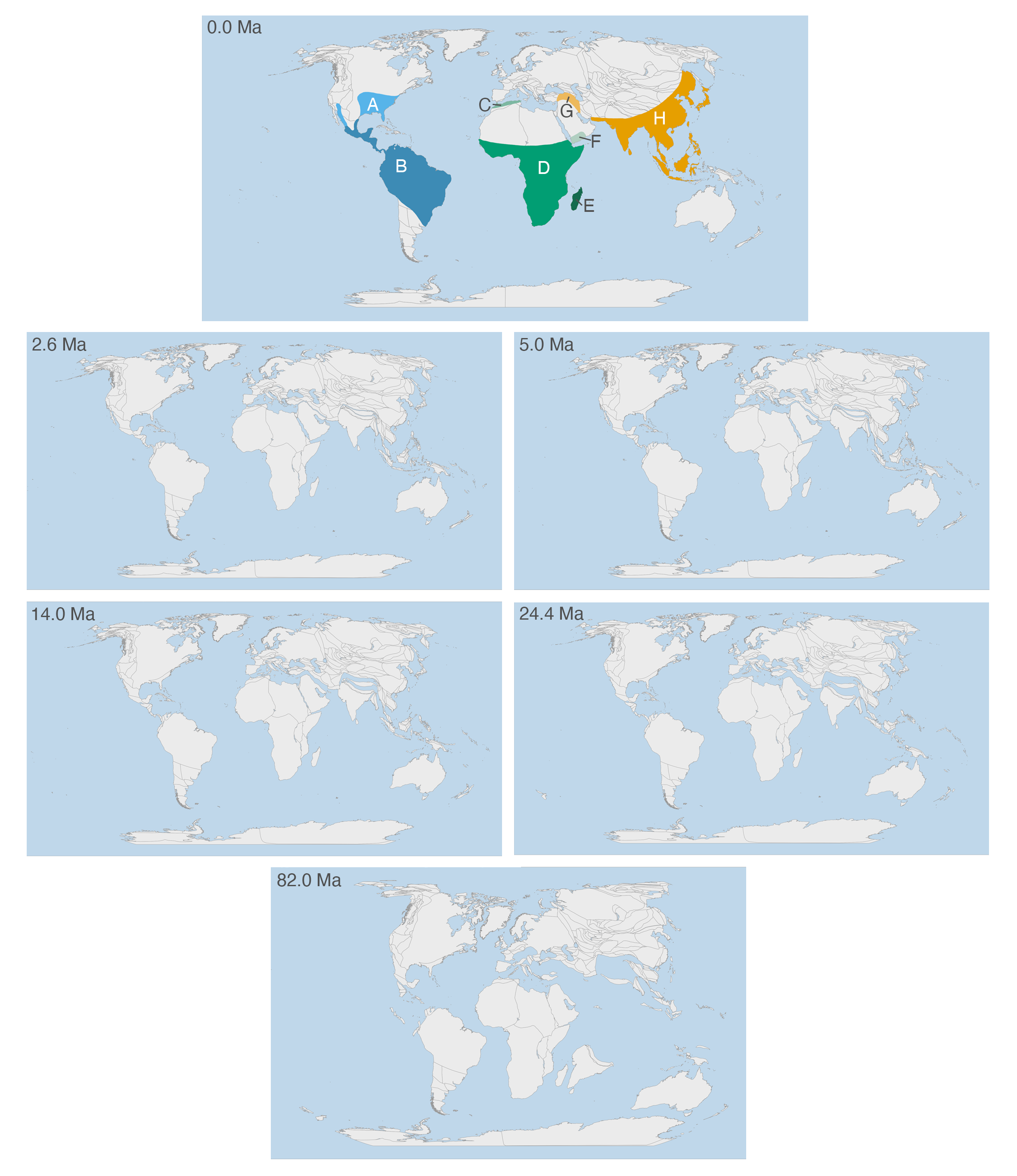
