## Supplementary material for "Long-distance dispersal drives global tropical distributions in a widespread moth lineage (Lepidoptera: Limacodidae)": Figure S6

**Fig. S6a: DEC**

ancstates: global optim, 2 areas max. d=0.0013; e=0.001; j=0; LnL=−76.80

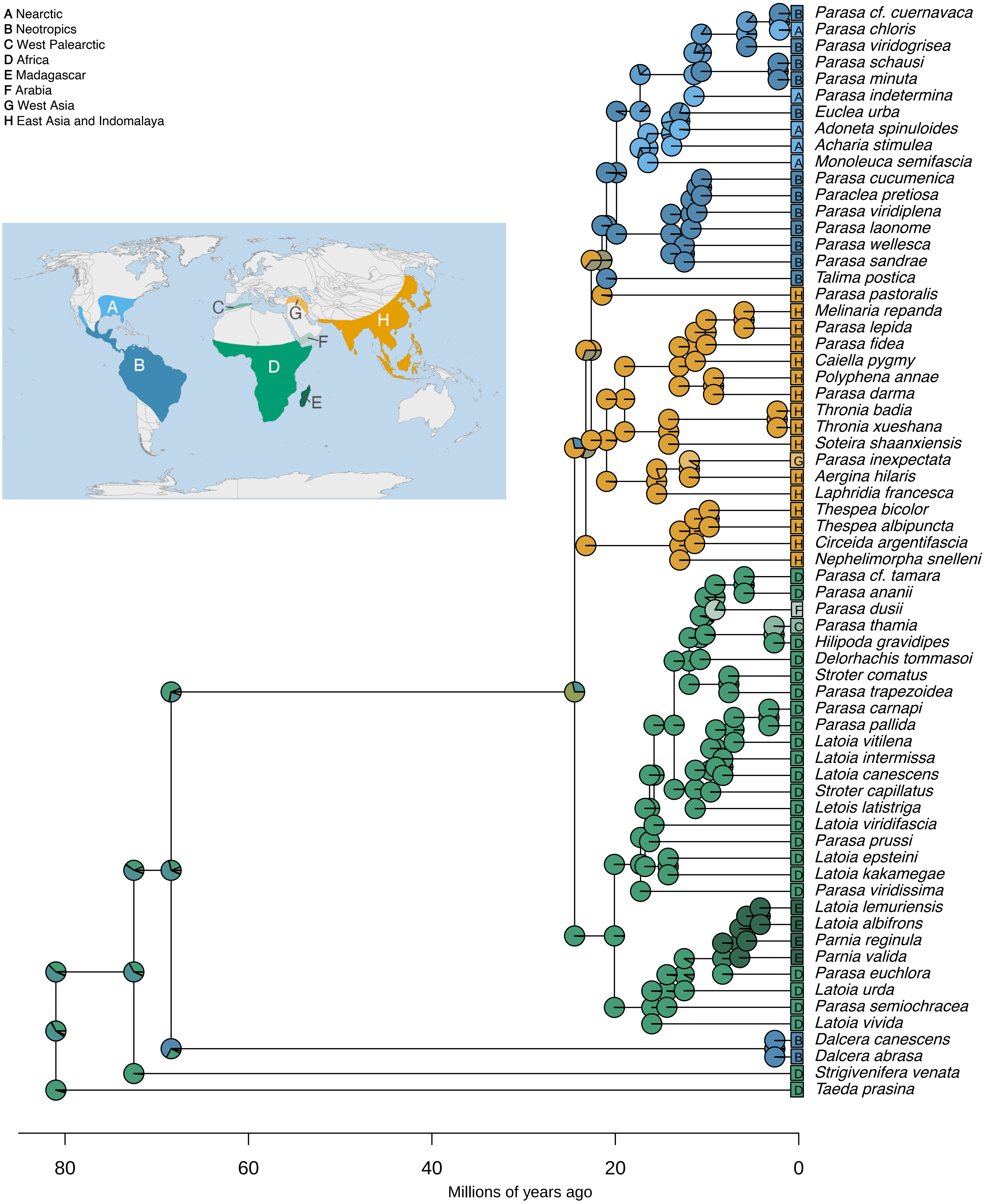

**Fig. S6b: DEC+J**

ancstates: global optim, 2 areas max. d=0; e=0; j=0.0145; LnL=−57.18

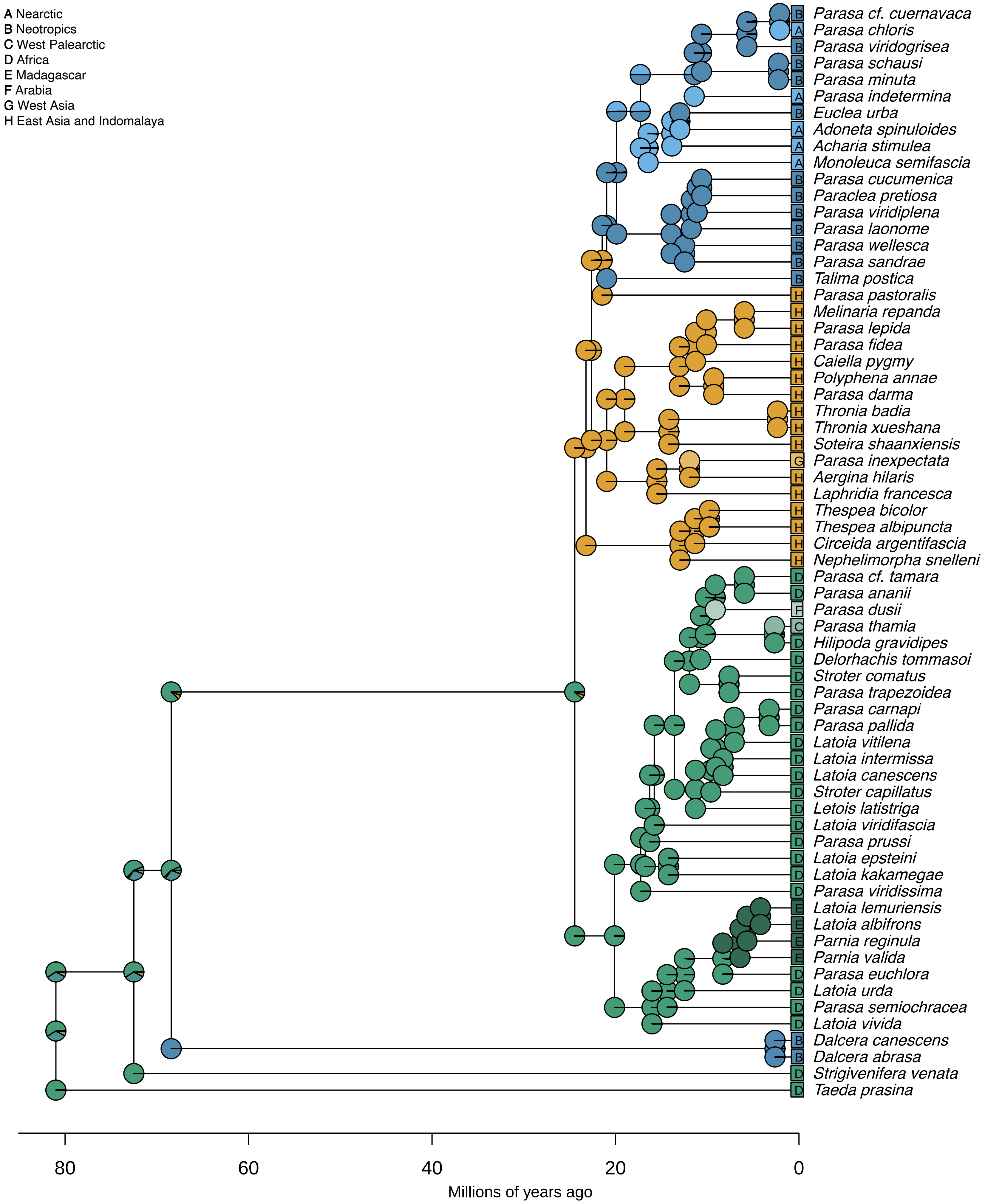

**Fig. S6c: DIVALIKE**

ancstates: global optim, 2 areas max. d=0.0016; e=9e−04; j=0; LnL=−75.04

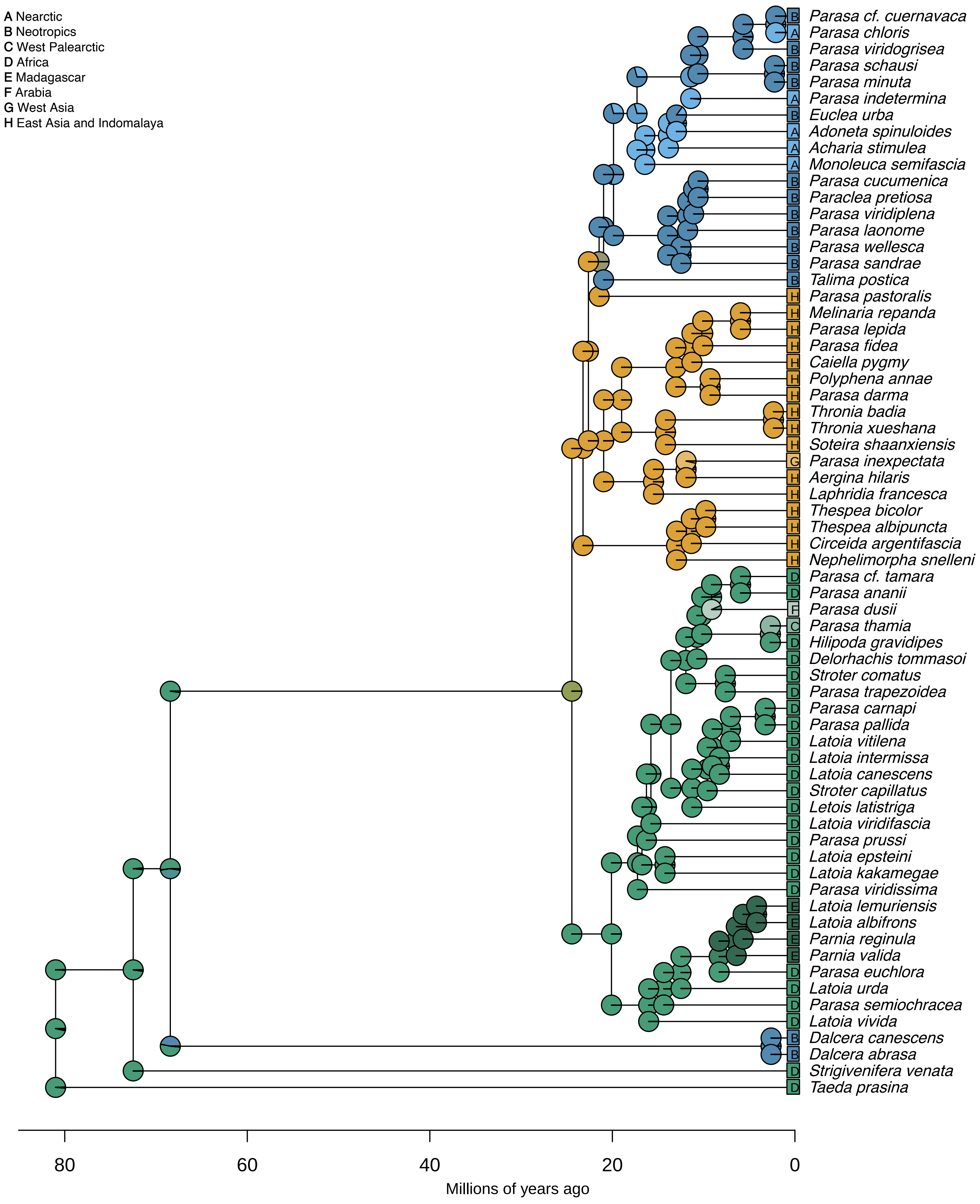

### Fig. S6d: DIVALIKE+J

ancstates: global optim, 2 areas max. d=0; e=0; j=0.0148; LnL=−57.68

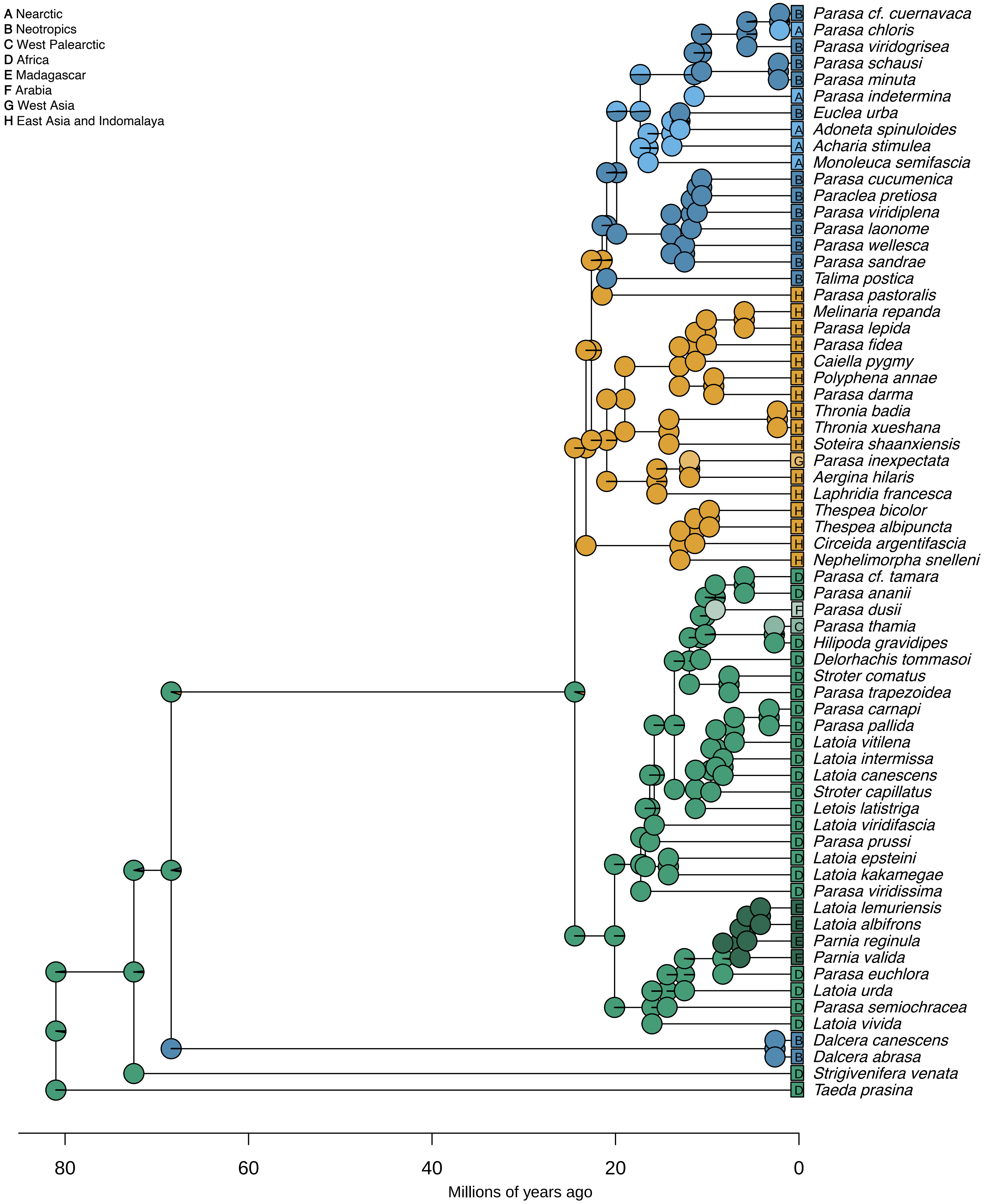

**Fig. S6e: BAYAREALIKE**

ancstates: global optim, 2 areas max. d=0.0015; e=0.0182; j=0; LnL=−103.07

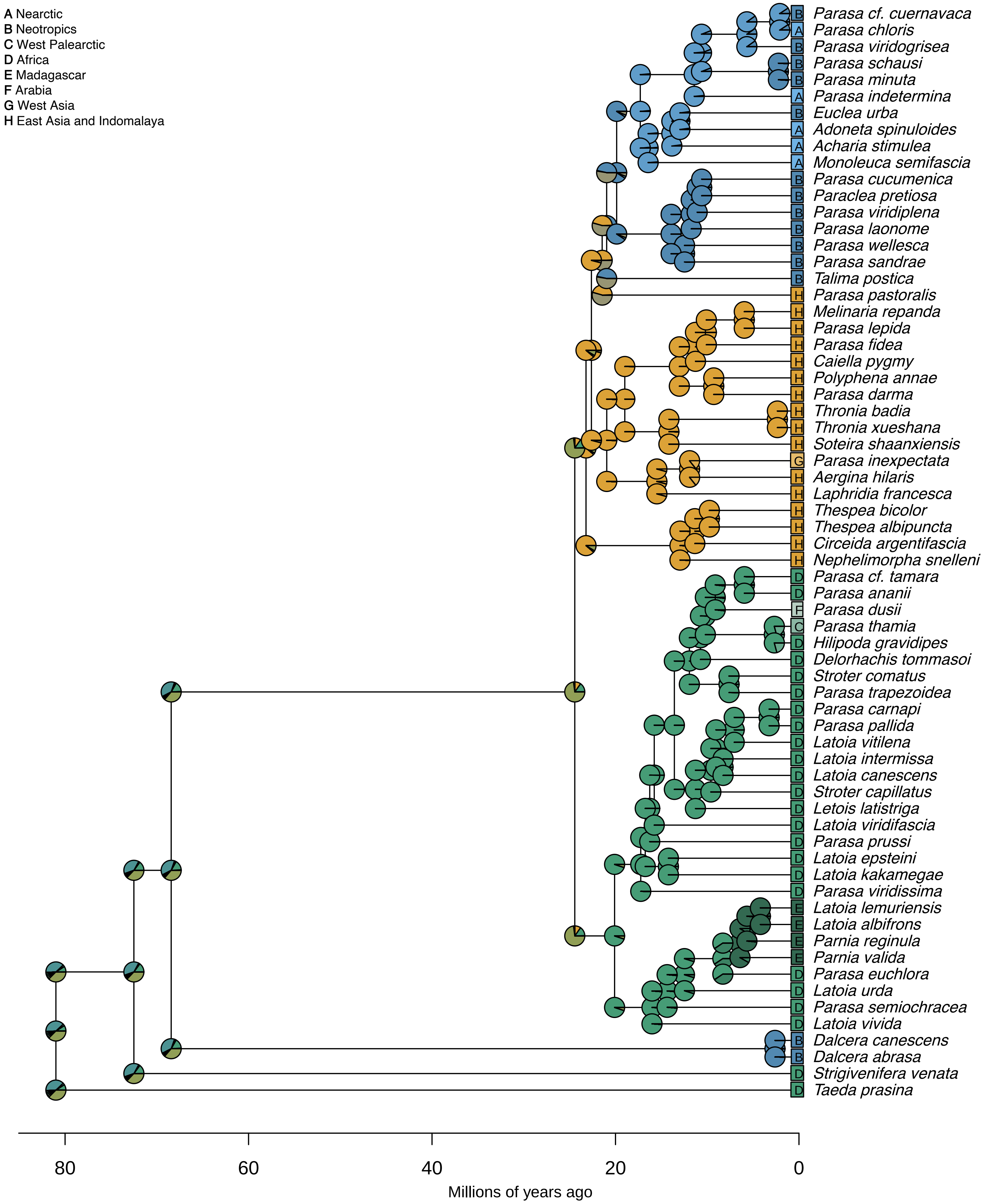

**Fig. S6f: BAYAREALIKE+J**

global optim, 2 areas max. d=0; e=0; j=0.0146; LnL=−57.71

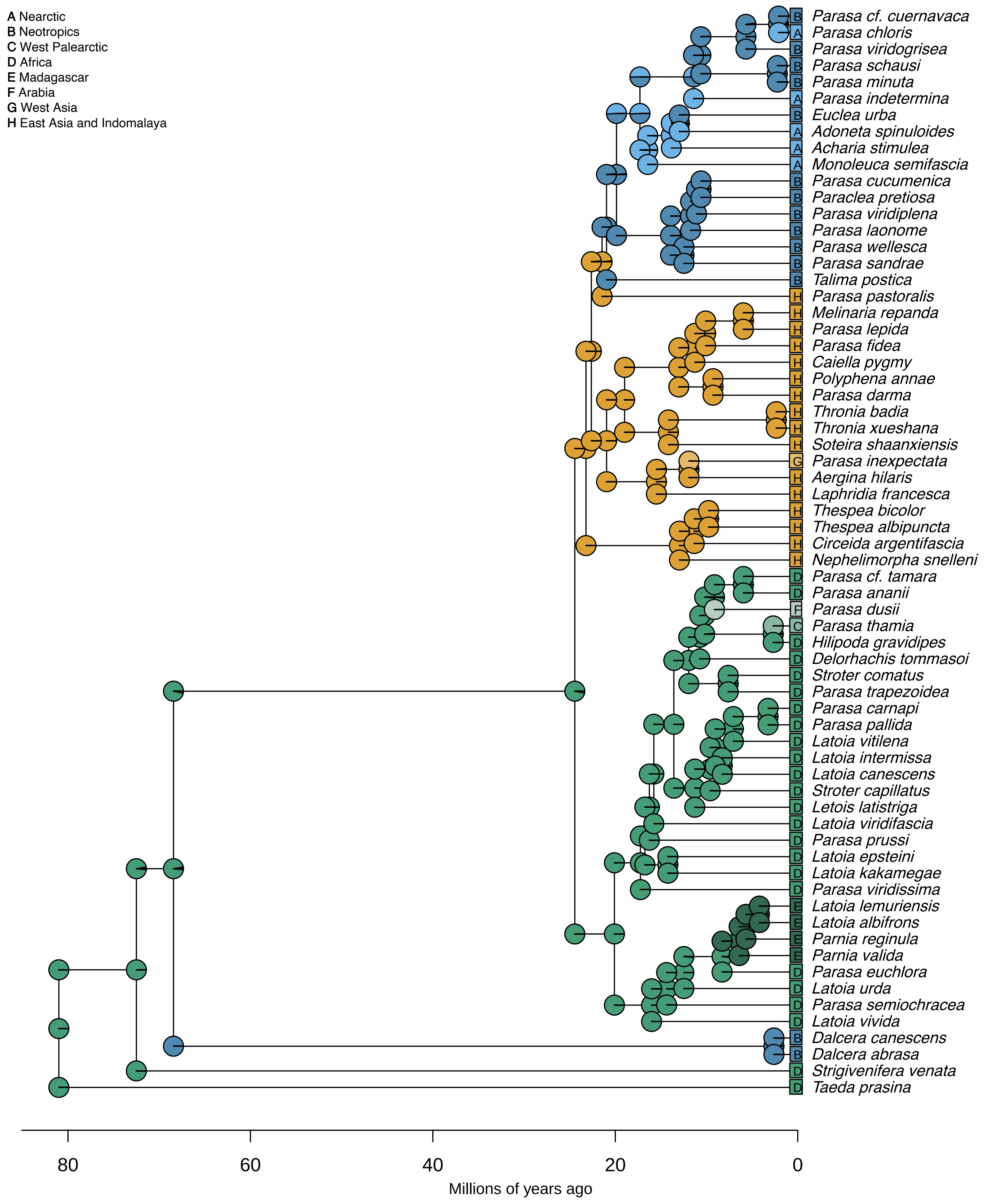

**Fig. S6g: Time-stratified DEC**

ancstates: global optim, 2 areas max. d=0.0036; e=6e−04; j=0; LnL=−74.37

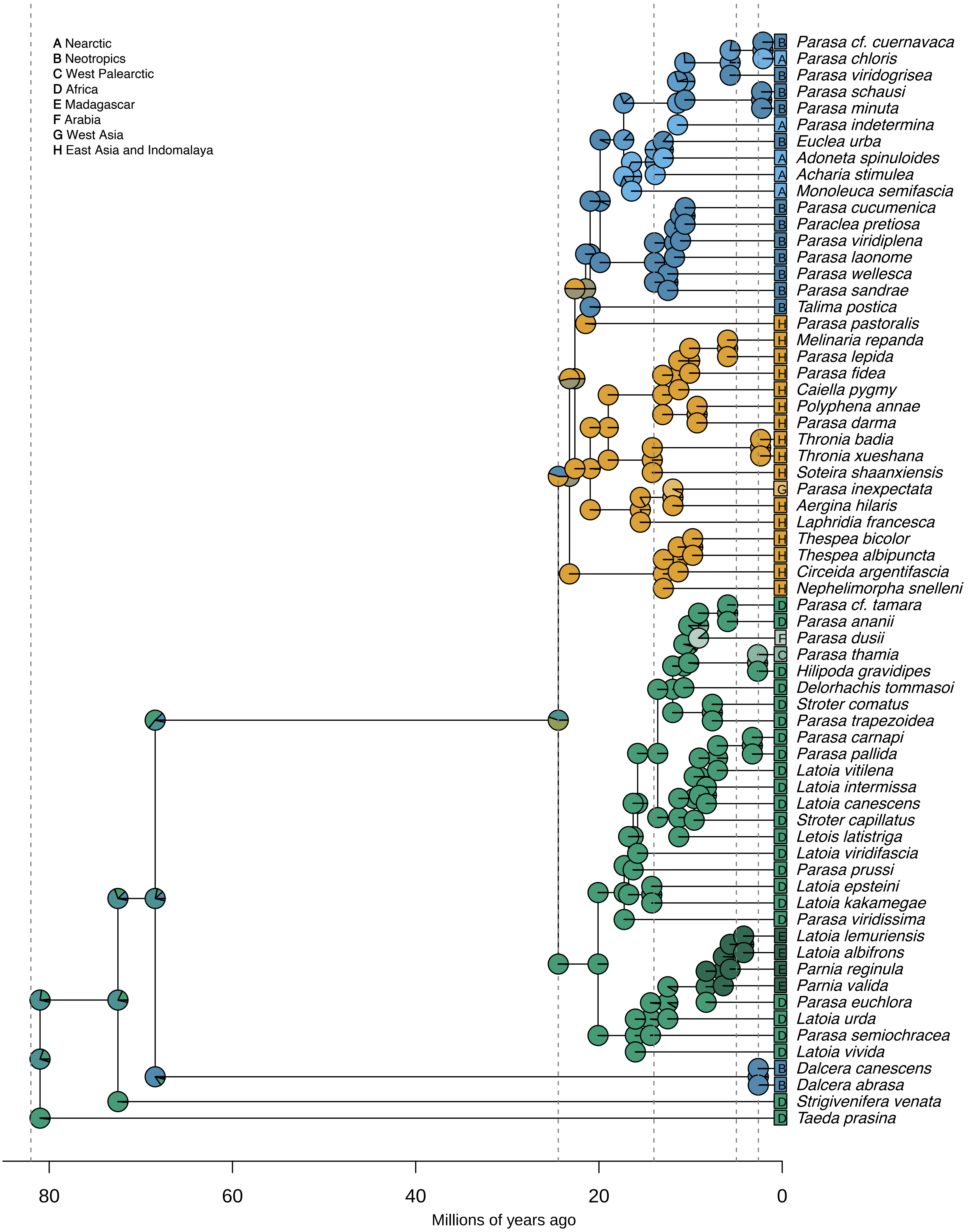

**Fig. S6h: Time-stratified DEC+J**

ancstates: global optim, 2 areas max. d=0; e=0; j=0.0446; LnL=−53.14

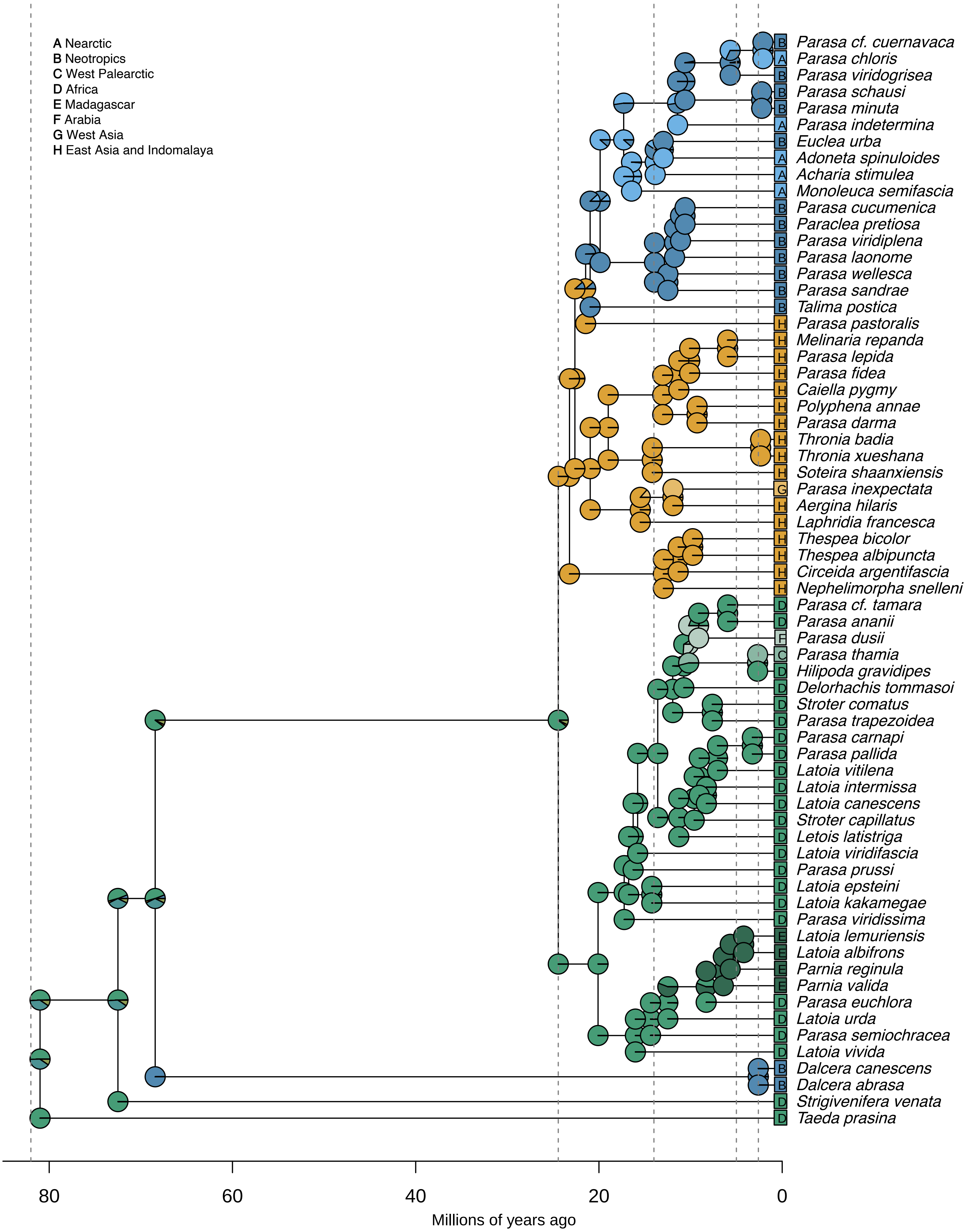

**Fig. S6i: Time-stratified DIVALIKE**

ancstates: global optim, 2 areas max. d=0.0047; e=9e−04; j=0; LnL=−71.98

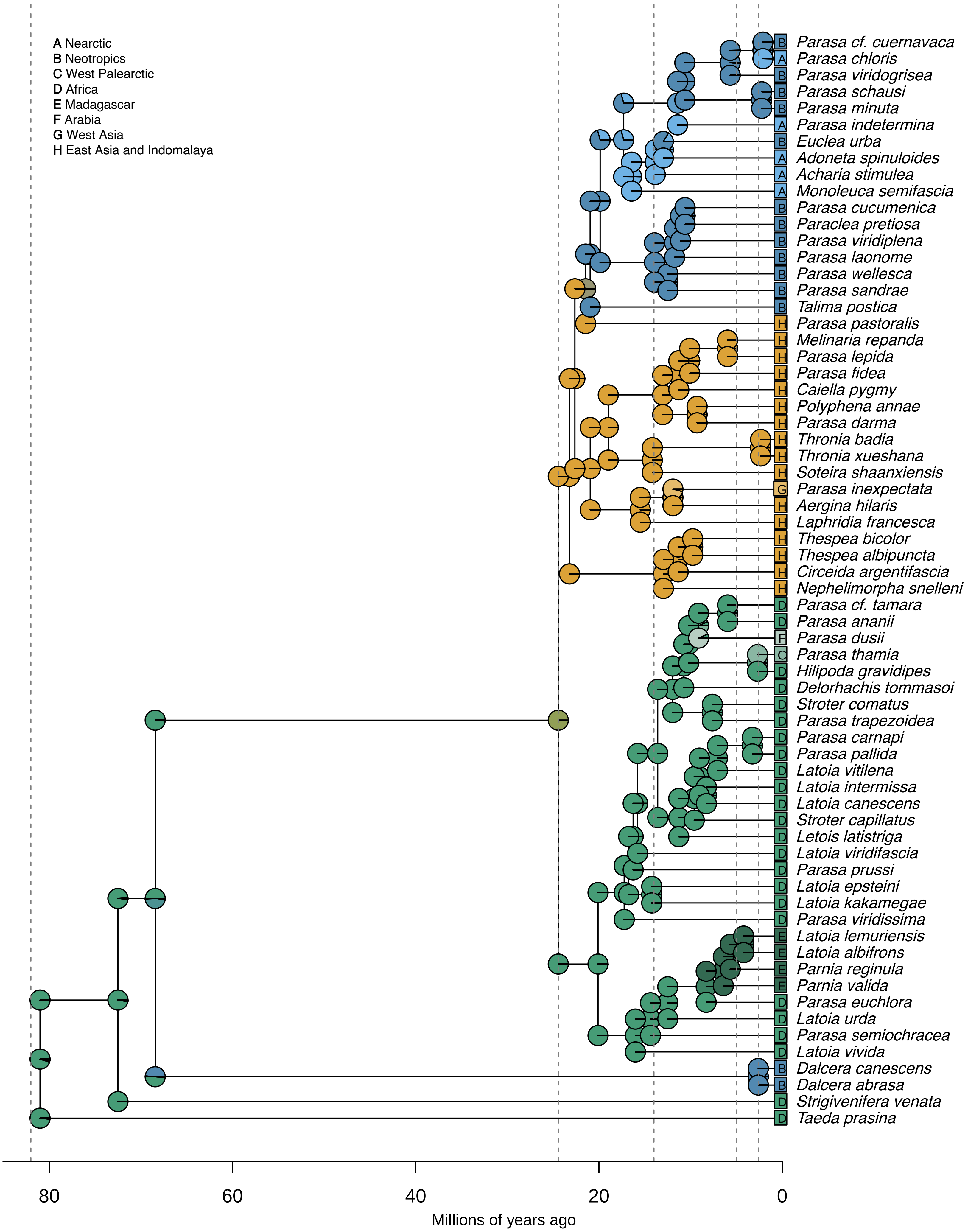

**Fig. S6j: Time-stratified DIVALIKE+J**

ancstates: global optim, 2 areas max. d=0; e=0; j=0.0454; LnL=–53.78

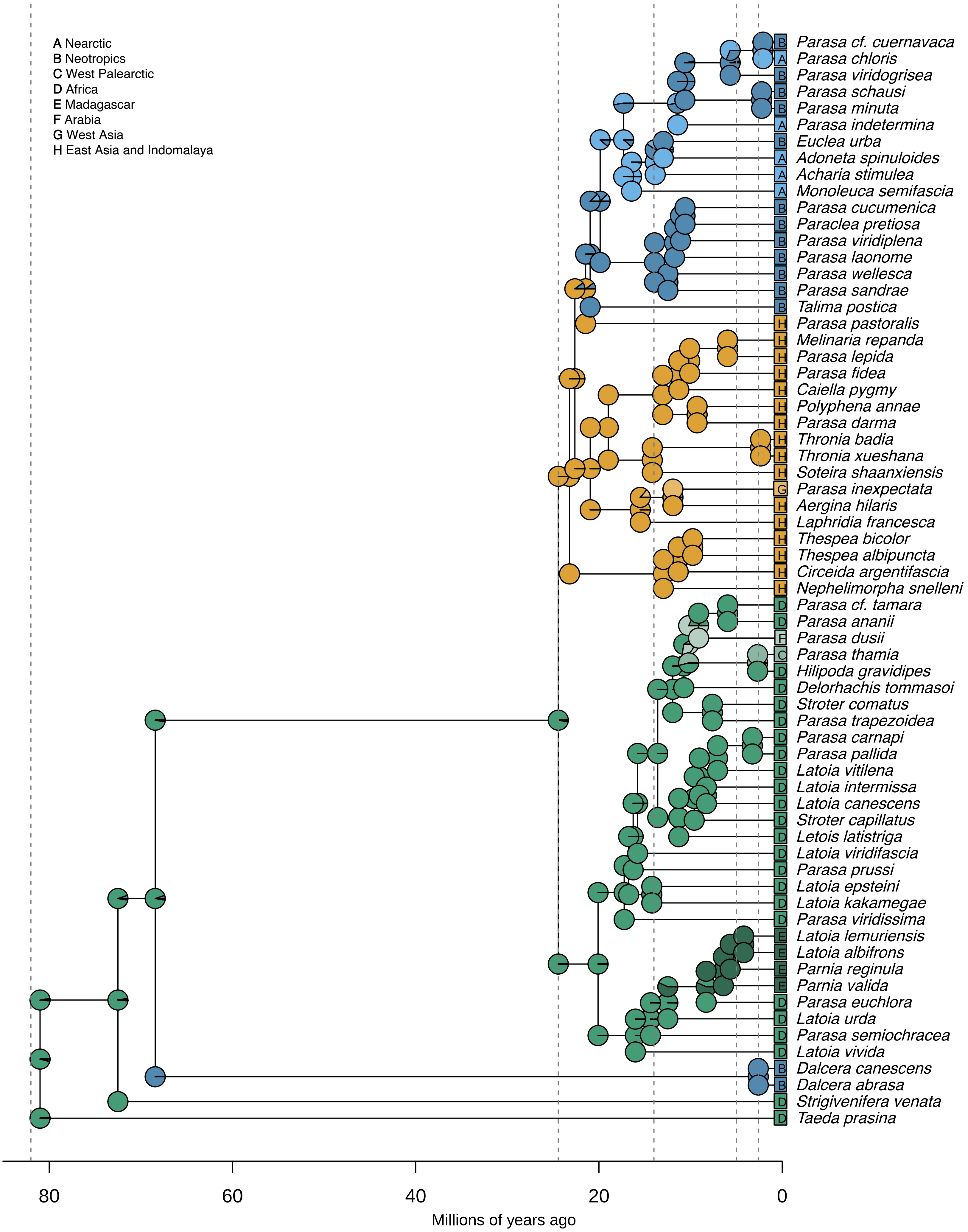

**Fig. S6k: Time-stratified BAYAREALIKE**

ancstates: global optim, 2 areas max. d=0.0045; e=0.018; j=0; LnL=−101.17

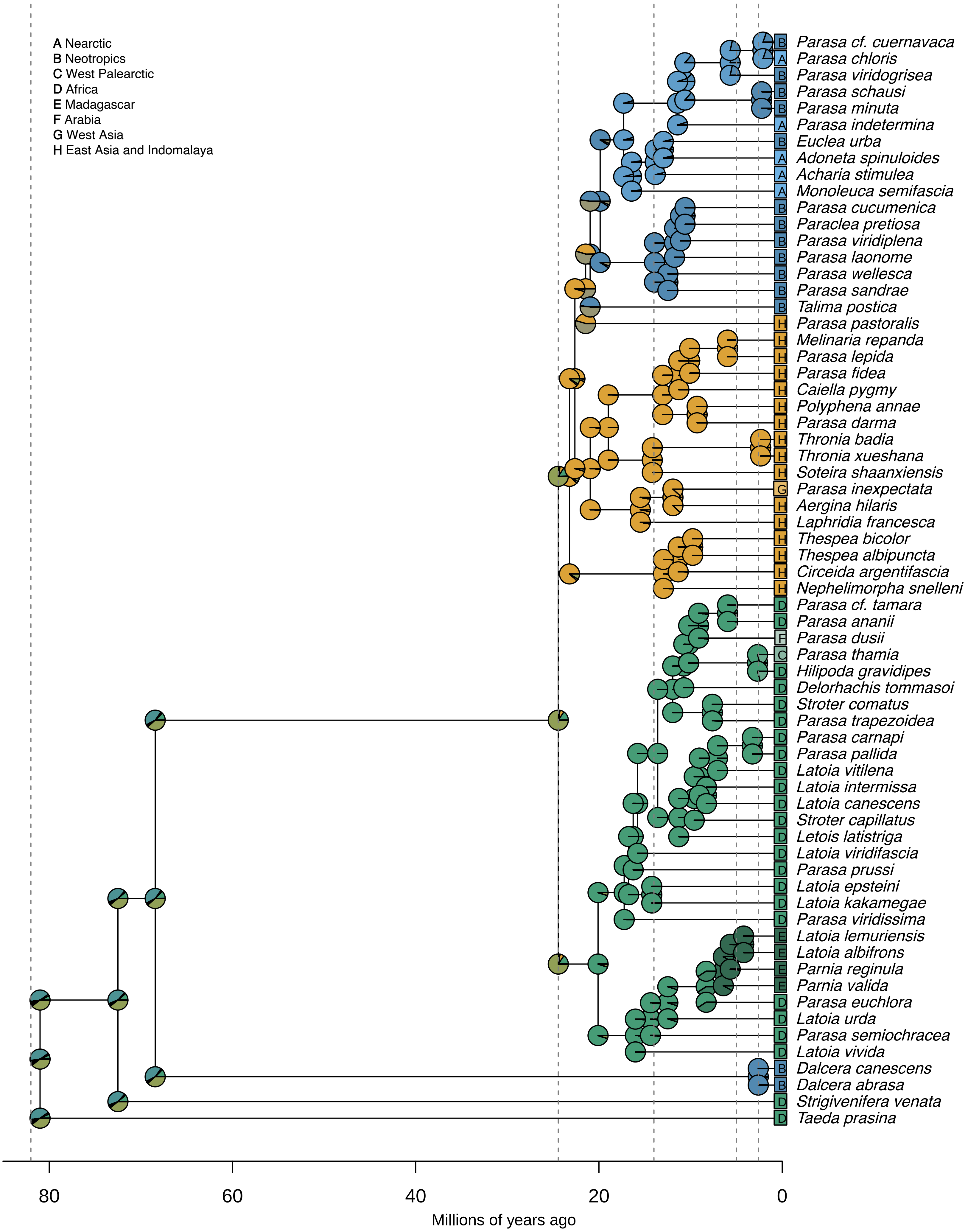

**Fig. S6I: Time-stratified BAYAREALIKE+J**

ancstates: global optim, 2 areas max. d=0; e=0; j=0.0442; LnL=−53.82

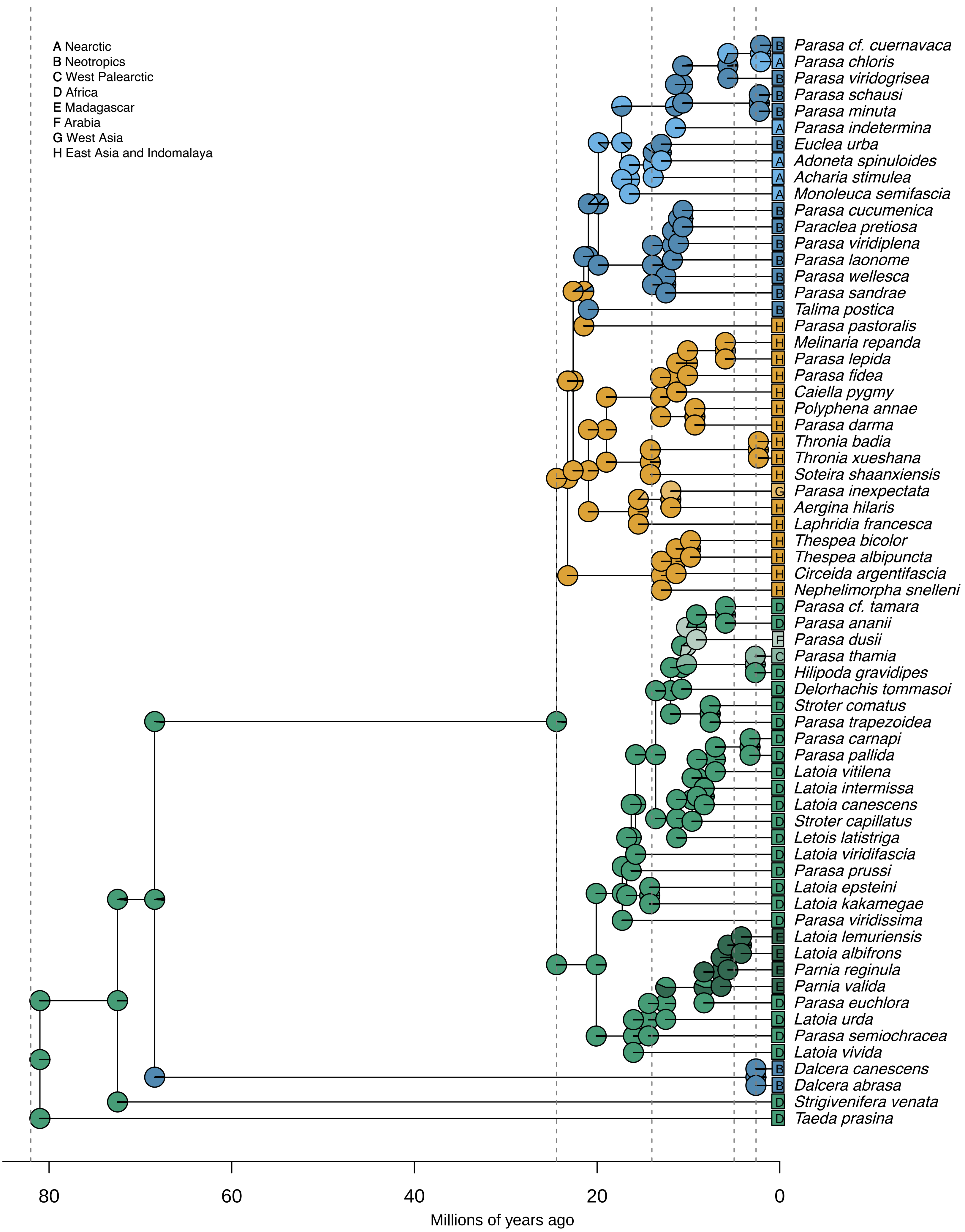
