## Supplementary file legends for "Long-distance dispersal drives global tropical distributions in a widespread moth lineage (Lepidoptera: Limacodidae)"

**Figures**

**Figure S1:** BUSCO gene recovery per sample, showing percentage recovered out of a possible 5,286 genes in the OrthoDB v.10 (Kriventseva *et al*., 2019) (*lepidoptera_odb10*) dataset.

**Figure S2:** Phylogenetic trees of the global *Parasa*-complex using ASTRAL-IV (within package ASTER; Zhang *et al.,* 2025) to combine 2,024 individual BUSCO gene trees (left) and IQ-TREE v2.2.2.6 (Minh *et al*., 2020) on a concatenated alignment totalling 3,229,490 bp of 2,024 BUSCO genes (right). Trees were first individually inspected in FigTree v1.4.4 (http://tree.bio.ed.ac.uk/software/figtree/), then tanglegrams were constructed with *cophylo* command in the package *phylogram* (Wilkinson & Davy, 2018) in R v4.3.2 (R Core Team 2023). All trees were initially rooted on the outgroup *Taeda prasina*, and then ladderised in decreasing node order with *ladderize* in the package *ape* (Paradis & Schliep, 2019). Branches/nodes were then not rotated (*rotate*=FALSE) when plotting with *cophylo* to ensure continuity of topological shape throughout. Support values shown with colour-coded key with the following abbreviations: local posterior probability (LPP) (for ASTRAL phylogeny, left), Shimodaira-Hasegawa-like approximate likelihood ratio test (SH-aLRT), ultrafast bootstrap (uBS) (for IQ-TREE phylogeny, right). a. Phylogenetic trees without sample *Ximacodes affinis*. b. Phylogenetic trees with sample *Ximacodes affinis.*

**Figure S3:** Time calibrated phylogeny of the *Parasa*-complex created with LSD2 v2.4.1 (<https://github.com/tothuhien/lsd2>) using the IQ-TREE phylogeny as input. Calibration points are represented with pink circles, and confidence intervals are represented with turquoise bars. Confidence intervals were generated by simulating 100 bootstrap trees in IQ-TREE.

**Figure S4:** Time calibrated phylogeny of the *Parasa*-complex created with treePL (Smith & O’Meara, 2012) using the IQ-TREE phylogeny as input. Calibration points are represented with pink circles, and confidence intervals are represented with turquoise bars. Confidence intervals were generated by simulating 100 bootstrap trees in IQ-TREE, then running treePL on each bootstrap replicates.

**Figure S5:** Geological landmass reconstructions created using the R package *rgplates* v0.6.1 (Kocsis *et al*., 2024) under the plate rotation model of Müller *et al.* (2019), based on periods specified for the time-stratified ancestral area estimation analyses.

**Figure S6:** Inferences of ancestral range reconstruction from all *BioGeoBEARS* (Matzke, 2013) models. a. Unconstrained DEC; b. Unconstrained DEC+*j*; c. Unconstrained DIVALIKE; d. Unconstrained DIVALIKE*+j*; e. Unconstrained BAYAREALIKE; f. Unconstrained BAYAREALIKE+*j*; g. Time-stratified DEC; h. Time-stratified DEC+*j*; i. Time-stratified DIVALIKE; j. Time-stratified DIVALIKE*+j*; k. Time-stratified BAYAREALIKE; l. Time-stratified BAYAREALIKE+*j*.

**Tables**

**Table S1:** Full details of species and their representative specimens sequenced for this study of the evolutionary history of the *Parasa*-complex. Table includes species name and region, specimen origin, storage method, DNA extraction yield, sequencing information, *de novo* assembly data and BUSCO gene recovery.

**Table S2:** Summary statistics for each of the 2,024 gene alignment files as created using AMAS (Borowiec, 2016) which remained after quality filtering.

**Table S3:** Results of quartet concordance, gene and site concordance calculations of the IQ-TREE phylogeny and ASTRAL phylogeny without *Ximacodes affinis*.

**Table S4:** Results of quartet concordance, gene and site concordance calculations of the IQ-TREE phylogeny and ASTRAL phylogeny including the sample of *Ximacodes affinis*.

**Table S5:** Dispersal rate matrices between all subregions for time-stratified *BioGeoBEARS* ancestral range reconstruction analyses, formulated based on observed distances between landmasses in geological reconstructions using the R package *rgplates* v0.6.1 (Kocsis *et al*., 2024) under the plate rotation model of Müller *et al.* (2019) (Fig. S5).

**Table S6:** Results of model selection (evaluated with log-likelihood values and AICc scores) for all unconstrained and time-stratified *BioGeoBEARS* ancestral range reconstruction analyses, with and without the jump parameter for founder events (+*j*; Matzke, 2014).

**Table S7:** Results of 1000 biogeographical stochastic mapping simulations in *BioGeoBEARS* under the DEC+*j* model to calculate the event frequencies of immigration and emigration between all subregions.
